# Heterozygous *Kmt2d* loss diminishes enhancers to render medulloblastoma cells vulnerable to combinatory inhibition of lysine demethylation and oxidative phosphorylation

**DOI:** 10.1101/2023.10.29.564587

**Authors:** Shilpa S. Dhar, Calena Brown, Ali Rizvi, Lauren Reed, Sivareddy Kotla, Constantin Zod, Janak Abraham, Jun-Ichi Abe, Veena Rajaram, Kaifu Chen, Min Gyu Lee

## Abstract

The histone H3 lysine 4 (H3K4) methyltransferase KMT2D (also called MLL4) is one of the most frequently mutated epigenetic modifiers in medulloblastoma (MB) and many other types of cancer. Notably, heterozygous loss of *KMT2D* is prevalent in MB and other cancer types. However, what role heterozygous *KMT2D* loss plays in tumorigenesis has not been well characterized. Here, we show that heterozygous *Kmt2d* loss highly promotes MB driven by heterozygous loss of the MB suppressor gene *Ptch* in mice. Heterozygous *Kmt2d* loss upregulated tumor-promoting programs, including oxidative phosphorylation and G-protein-coupled receptor signaling, in *Ptch^+/−^-*driven MB genesis. Mechanistically, both downregulation of the transcription-repressive tumor suppressor gene NCOR2 by heterozygous *Kmt2d* loss and upregulation of the oncogene *MycN* by heterozygous *Ptch* loss increased the expression of tumor-promoting genes. Moreover, heterozygous *Kmt2d* loss extensively diminished enhancer signals (e.g., H3K27ac) and H3K4me3 signature, including those for tumor suppressor genes (e.g., *Ncor2*). Combinatory pharmacological inhibition of oxidative phosphorylation and the enhancer-decommissioning H3K4 demethylase LSD1 drastically reduced tumorigenicity of MB cells bearing heterozygous *Kmt2d* loss. These findings reveal the mechanistic basis underlying the MB-promoting effect of heterozygous *KMT2D* loss, provide a rationale for a therapeutic strategy for treatment of KMT2D-deficient MB, and have mechanistic implications for the molecular pathogenesis of other types of cancer bearing heterozygous *KMT2D* loss.

## INTRODUCTION

Histone lysine (K) methylation is a central player for epigenetic and transcriptional regulation of gene expression. This modification is associated with activation or silencing of gene expression, depending on the site of lysine methylation. For example, methylation at H3K4 is generally associated with gene activation, whereas methylation at H3K27 is linked to gene repression. Histone lysine methylation is catalyzed by lysine methyltransferases ^1,2^ but can be reversed by lysine demethylases ^3–5^. Interestingly, sequencing studies of cancer genomes have shown that these modifiers are often mutated in cancer. In particular, the histone methyltransferase KMT2D (also called MLL4, MLL2, and ALR) is one of the most frequently mutated histone modifiers in cancer ^6,7^.

KMT2D is a member of the SET/COMPASS (COMplex of Proteins ASsociated with Set1) family of histone H3K4 methyltransferases ^8^. Notably, heterozygous loss (single-allelic loss) of *KMT2D* is prevalent in MB and other types of cancer according to datasets in the cBioportal (**Extended Data Fig. 1a**). The effects of tissue-specific homozygous loss of *Kmt2d* on tumorigenesis in various tissues have been studied ^9–14^. However, what role heterozygous *KMT2D* loss plays in tumorigenesis **remains not well characterized**.

MB is one of the most common childhood brain tumors (up to 20% of all childhood brain tumors) and is located in the cerebellum ― a brain region that is central for controlling motor coordination and balance ^15^. MB is classified as a type of high-grade malignant primary tumors ^16^. MB can be categorized into four major molecular subgroups: sonic hedgehog (SHH), wingless (WNT), Group 3, and Group 4 ^17,18^. The 5-year survival rates of SHH, Group 3, and Group 4 are 50%–75% whereas that of SHH subgroup (10% of MBs) is ∼95% ^19^. MB development is often associated with altered signaling pathways, such as altered SHH and WNT pathways ^18^. For example, *PTCH* (also called *PTCH1*) is the most frequently mutated gene in the SHH pathway. Overall, the mutation rate of *KMT2D* in MB is 8%‒11%, which is similar to that of *PTCH* in MB ^19–24^. KMT2D mutations occur in all four subgroups. Most of the *KMT2D* mutations in MB are truncations according to the datasets in the cBioPortal. Besides its mutations, KMT2D is downregulated in another significant portion (∼ up to 5%) of MBs (**Extended Data Fig. 1b**). These KMT2D alterations (up to 15.1%), which generally result in KMT2D deficiency, correlate with poor survival in MB patients (**Extended Data Fig. 1c**), consistent with a previous report showing that low KMT2D levels were associated with poor survival in MB patients ^25^.

Our previous study using *Nestin-Cre Kmt2d*^fl/fl^ mice showed that brain-wide homozygous loss of *Kmt2d* induced MB in 7-month-old adult mice, indicating that KMT2D is a critical MB suppressor ^9^. Considering that MB occurs predominantly in children and adolescents younger than age 15 years, this finding raises the possibility that *KMT2D* loss needs to cooperate with other genetic or epigenetic oncogenic event(s) to induce early development of MB. In addition, as the vast majority of the *KMT2D* mutations in human MB were heterozygous ^25^, it is plausible that heterozygous *KMT2D* loss would be important for MB genesis. In fact, heterozygous loss of critical tumor suppressor genes contributes to tumorigenesis. For example, heterozygous inactivation of PTEN results in haploinsufficiency for tumorigenesis ^26^, and heterozygous p53 loss causes haploinsufficiency for multiple myeloma ^27^. For these reasons, we sought to assess whether and how *Kmt2d* loss, including heterozygous *Kmt2d* loss, cooperates with other oncogenic event to promote MB genesis.

## RESULTS

### *Nestin-Cre*-mediated heterozygous loss of *Kmt2d* in the brain accelerates *Ptch^+/−^*-driven MB genesis in mice

To test whether *Kmt2d* loss cooperates with other MB-relevant oncogenic event(s) to induce MB, we sought to assess the effect of *Kmt2d* loss and other MB-relevant gene alteration on MB genesis using mouse models. For other MB-relevant gene alteration, we chose *Ptch^+/−^* mice for the following reasons: (**i**) PTCH is a critical MB suppressor ^28^; (**ii**) *Ptch^+/−^* mice (i.e., mice with heterozygous *Ptch* loss) are a well characterized MB model with 10%–20% MB incidence within postnatal 10–25 weeks ^28–30^; (**iii**) *PTCH* is one of the most frequently mutated tumor suppressor genes in MB ^20^; and (**iv**) *KMT2D* mutations has a better tendency to co-occur with *PTCH* mutations than with several other mutations in MB samples ^23,31,32^.

We generated genetically engineered mouse models (GEMMs), including *Nestin*-*Cre Kmt2d*^fl/+^ *Ptch*^+/−^ (referred to as <u>NKP</u>) and *Nestin-Cre Kmt2d*^fl/fl^ *Ptch*^+/−^*. Nestin-Cre* deletes a floxed gene in many brain regions because *Nestin-Cre*^+/−^ mice express Cre recombinase (as early as E8.5) under the *Nestin* promoter and enhancer that drive gene expression mainly in neural stem cells and neural progenitor cells in the central nervous system ^33^. *Nestin-Cre Kmt2d*^fl/fl^ *Ptch^+/−^* mice could not be monitored for MB genesis because they died by postnatal day 15 (P15). Interestingly, magnetic resonance imaging (MRI) of NKP mice showed that *Nestin-Cre*-mediated heterozygous loss of *Kmt2d* drastically enhanced *Ptch^+/−^-*driven MB genesis in the cerebellum (**Fig. 1a**). H & E staining showed that MBs were visible in 4-month-old NKP cerebella (**Fig. 1b**). However, MBs were not visible in 2-month-old NKP cerebella (**Extended Data Fig. 2a,b**). These tumors were classic MBs, as the tumor cells were small, round, and undifferentiated and had a similar size of nuclei without an obvious pleomorphism (**Fig. 1b**). Average tumor volume was much bigger in NKP mice than in *Ptch*^+/−^ mice (**Fig. 1c**). In line with this, 52% of NKP mice (13/25) developed MBs whereas 14% of *Ptch*^+/−^ mice (2/14) did (**Fig. 1d**), and NKP mice showed lower survival than did *Ptch*^+/−^ mice and *Nestin-Cre Kmt2d*^fl/fl^ mice (**Fig. 1e**). As these results showed that heterozygous *Kmt2d* loss alone did not induce MB but highly promoted *Ptch*^+/−^-induced MB genesis, this indicates that heterozygous *Kmt2d* loss cooperates with heterozygous *Ptch* loss for MB genesis.

**Figure 1:**
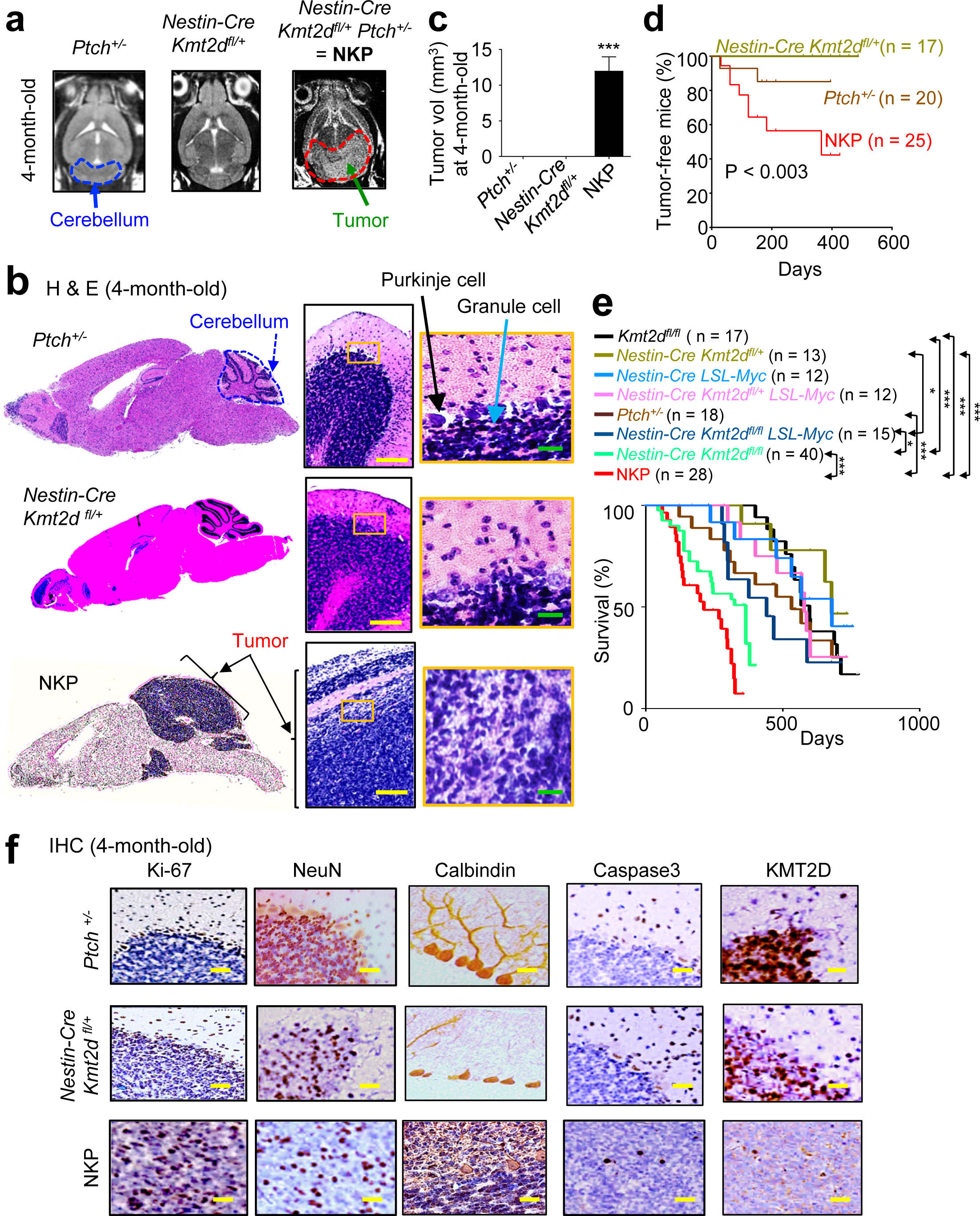
*Nestin-Cre*-mediated heterozygous loss of *Kmt2d* promotes *Ptch*-driven MB genesis in mice. (**<u>a</u>**) Horizontal magnetic resonance images (MRI) of *Ptch^+/−^, Nestin-Cre Kmt2d^fl/+^*, and *Nestin-Cre Kmt2d^fl/+^ Ptch^+/−^* (NKP) mice (4-month-old). The blue dotted line shows normal cerebellum, and the red dotted line denotes tumors in the cerebellum in NKP mice. (**<u>b</u>**) H&E staining of sagittal brain sections of *Ptch^+/−^, Nestin-Cre Kmt2d^fl/+^,* and NKP mice. Whole brains and cerebellar regions are shown. The yellow-boxed regions are presented at higher magnification. Yellow scale bars, 200 μm; green scale bars, 40 μm. (**<u>c</u>**) Quantification of tumor volumes in *Ptch^+/−^, Nestin-Cre Kmt2d^fl/+^*, and NKP cerebella using MRI data. Data are presented as the mean ± SEM (error bars) (n = 3). (**<u>d</u>**) MB-free percentages of *Ptch^+/−^, Nestin-Cre Kmt2d^fl/+^*, and NKP mice. (**<u>e</u>**) Kaplan-Meier survival analysis of different mouse groups using a log-rank test. * p<0.05, ** p<0.01, *** p<0.001. (**f**) Immunohistochemistry (IHC) analysis for Ki-67, NeuN, Calbindin, Caspase3, and KMT2D in *Ptch^+/−^, Nestin-Cre Kmt2d^fl/+^*, and NKP tumor tissues. See also **Extended Data Fig. 1–4**.

We examined whether the status of heterozygous *Kmt2d* loss is maintained during MB genesis. Our results showed that the *Kmt2d* gene in the wild type (WT) allele in NKP tumors was likely intact (**Extended Data Fig. 2c**). This indicates that loss of heterozygosity of *Kmt2d* (i.e., loss of non-floxed *Kmt2d* gene) was unlikely to occur in NKP tumors and confirms the MB-promoting effect of heterozygous *Kmt2d* loss.

We also sought to assess the effect of *Kmt2d* loss on another MB-relevant oncogenic event in addition to heterozygous *Ptch* loss. MYC amplification frequently occurs in Group 3 MB. Interestingly, p53 alterations (p53 loss or a dominant negative form of p53), along with MYC overexpression, induced Group 3-like MB although p53 alterations or MYC overexpression alone do not induce MB ^34–37^. Our previous study using *Nestin-Cre Kmt2d*^fl/fl^ mice showed that brain-specific *Kmt2d* loss induced MBs that are positive for Group 3 MB markers ^9^. This led us to determine whether *Kmt2d* loss similar to p53 loss cooperates with MYC overexpression for MB genesis. We generated *Nestin-Cre LSL-Myc Kmt2d*^fl/+^ and *Nestin-Cre Kmt2d*^fl/fl^ *LSL-Myc* mice. *Nestin-Cre* induced *Myc* overexpression via Cre-mediated deletion of the LSL (loxP-STOP-loxP) cassette from *LSL-Myc* locus (**Extended Data Fig. 3a**). However, Myc overexpression did not cooperate with homozygous or heterozygous loss of *Kmt2d* and appeared to negatively impact *Kmt2d*-loss–driven MB genesis in *Nestin-Cre Kmt2d*^fl/fl^ mice (**Fig. 1e and Extended Data Fig. 3a,b**). This result can be explained by the previous report that MYC overexpression caused Myc-induced cellular apoptosis that can be mediated by p53 while not inducing MB ^34^. In line with this report, staining of cleaved caspase 3 were higher in MYC-overexpressed cerebella than in *Nestin-Cre Kmt2d*^fl/fl^ cerebella, indicating that apoptosis occurred in MYC-overexpressed cerebella (**Extended Data Fig. 3a**). These results indicate that KMT2D unlike p53 may not regulate apoptosis. For these reasons, we focused on understanding the role of heterozygous *Kmt2d* loss in accelerating *Ptch^+/−^-*driven MB genesis.

### *Nestin-Cre*-mediated heterozygous loss of *Kmt2d* increases cell proliferation in the *Ptch^+/−^* cerebellum and aberrantly impacts cerebellar neurons

Because tumorigenesis is promoted by increased cell proliferation ^38^, we assessed whether heterozygous *Kmt2d* loss increases cell proliferation in the cerebellum. We compared the cellular states of NKP cerebella bearing MB with those of *Ptch^+/−^* and *Nestin-Cre Kmt2d*^fl/+^ cerebella (another control) using IHC staining of Ki-67 (a proliferation marker). Our results showed that Ki-67-positive cells were slightly increased in 2-month-old NKP cerebella and highly increased in 4-month-old NKP cerebella (**Fig. 1f and Extended Data Fig. 3c**), indicating that cells in 4-month-old NKP cerebella are not neuronal cells but highly proliferating tumor cells.

Purkinje cells and granule cells represent the major types of cerebellar neurons ^39^. Granule cells are the vast majority of neurons in the cerebellum and are glutamatergic excitatory neurons ^40^. NeuN heavily labels the granule cells (due to their small size and dense packing) in the cerebellum although it is a marker for most mature types of neuronal cells. Purkinje cells are a type of GABAergic inhibitory neurons that are responsible for main computations in the cerebellum. A well-known neuronal marker for Purkinje cells is Calbindin. As compared to *Ptch^+/−^* cerebella, NeuN and Calbindin staining was slightly decreased in 2-month-old NKP cerebella while being drastically reduced in 4-month-old NKP cerebella, suggesting that Purkinje cells and granule cells are negatively impacted by heterozygous *Kmt2d* loss (**Fig. 1f and Extended Data Fig. 3c**). To link NKP MBs to MB subgroups, we examined their gene expression profiles and performed immunofluorescence (IF) staining of 4-month-old *Ptch*^+/−^ and NKP cerebella using subtype-specific markers ^18^. Results showed that NKP MB was positive for the SHH marker GAB1 and the group 3 marker MYC and that expression of group 3- and SHH-relevant genes was enriched in NKP MB (**Extended Data Fig. 4a,b**), suggesting that the NKP MB would have characteristics of SHH and group 3.

### *Atoh1-Cre*-mediated heterozygous loss of *Kmt2d* greatly promotes *Ptch^+/−^*-driven MB genesis

Interestingly, KMT2D levels were higher in the cerebellum than in other brain regions (**Extended Data Fig. 5a,b**). *Atoh1*-*Cre* deletes a floxed gene largely in cerebellar granule cell (GC) lineage, which generates GCs ^40^. Cerebellar GC progenitors can serve as a cell of origin for MB ^41^. For these reasons, we generated *Atoh1*-*Cre Kmt2d*^fl/+^ *Ptch^+/−^* (referred to as <u>AKP</u>) mice using *Atoh1*-*Cre* and monitored the onset of MB by biweekly performing MRI (starting from 0.5‒ 1 month). Similar to *Nestin-Cre Kmt2d*^fl/fl^ *Ptch^+/−^* mice, *Atoh1*-*Cre Kmt2d*^fl/fl^ *Ptch^+/−^* mice died within P15. Remarkably, *Atoh1*-*Cre*-mediated heterozygous loss of *Kmt2d* highly increased *Ptch^+/−^*-driven MB in 2-month-old AKP mice (**Fig. 2a–d**) and as a consequence reduced mouse survival time (**Fig. 2e**). *Atoh1*-*Cre*-mediated heterozygous loss of *Kmt2d* appeared to unlikely induce loss of heterozygosity of *Kmt2d* (**Extended Data Fig. 5c**). In 2-month-old AKP cerebella compared to *Ptch^+/−^* cerebella, the number of Ki-67-positive cells were strongly increased, and NeuN and Calbindin staining was drastically decreased (**Fig. 2f**). These results indicate that MB cells proliferate for tumorigenesis in the AKP cerebella.

**Figure 2:**
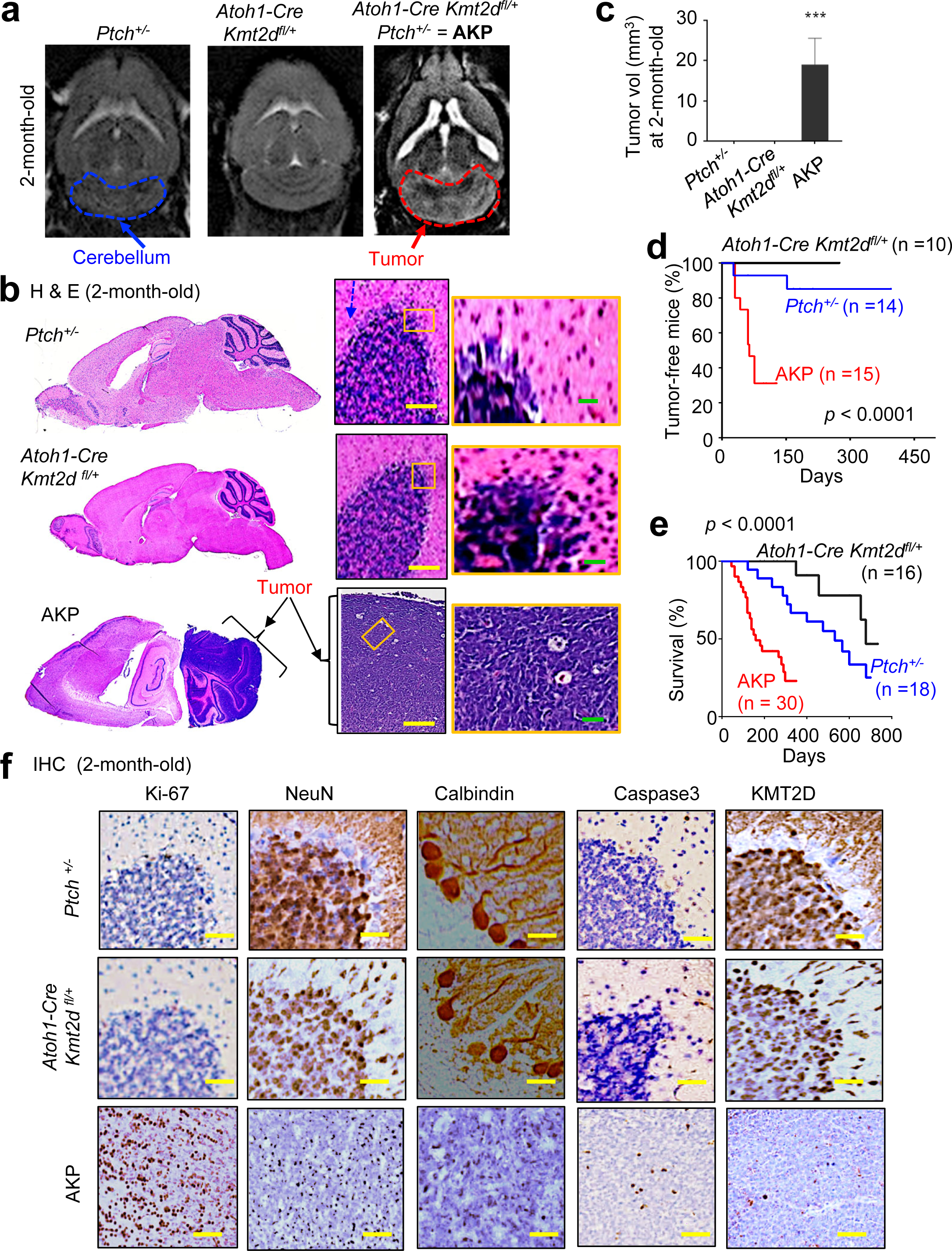
*Atoh1-Cre*-mediated heterozygous loss of *Kmt2d* highly increases *Ptch*-driven MB genesis in mice. (**<u>a</u>**) MRI images of *Ptch^+/−^, Atoh1-Cre Kmt2d^fl/+^*, and *Atoh1-Cre Kmt2d^fl/+^ Ptch^+/−^*(AKP) mice (2-month-old). The blue dotted line shows normal cerebellum, and the red dotted line denotes tumors in the cerebellum in AKP mice. (**<u>b</u>**) H&E staining of sagittal brain sections of *Ptch^+/−^, Atoh1-Cre Kmt2d^fl/+^*, and AKP mice. Whole brains and cerebellar regions are shown. The yellow-boxed regions are presented at higher magnification. Scale bars: 200 μm (yellow), 40 μm (green). (**<u>c</u>**) Quantification of tumor volumes in *Ptch^+/−^, Atoh1-Cre Kmt2d^fl/+^*, and AKP cerebella using MRI data. Data are presented as the mean ± SEM (error bars) (n = 3). (**<u>d</u>**) MB-free percentages in *Ptch^+/−^, Atoh1-Cre Kmt2d^fl/+^*, and AKP mice. (**<u>e</u>**) Kaplan-Meier survival analysis of different mouse groups using a log-rank test. (**<u>f</u>**) Immunohistochemistry (IHC) analysis for Ki-67, NeuN, Calbindin, Caspase3, and KMT2D in *Ptch^+/−^, Atoh1-Cre Kmt2d^fl/+^*, and AKP tumor tissues. See also **Extended Data Fig. 5**.

### Heterozygous *Kmt2d* loss upregulates tumor-promoting programs

To understand the molecular mechanism by which heterozygous *Kmt2d* loss promotes *Ptch*^+/−^ -driven MB genesis, we isolated the cerebella from *Ptch*^+/−^ and NKP (or AKP) mice bearing MB and then compared gene expression profiles between *Ptch*^+/−^ and NKP cerebella (or AKP cerebella) using RNA-seq. We found that heterozygous *Kmt2d* loss commonly upregulated almost identical set of tumor-promoting programs in both NKP and AKP cerebella compared to *Ptch*^+/−^ cerebella (**Fig. 3a,b**). Such tumor-promoting programs include oxidative phosphorylation (OXPHOS) pathway, oxidation-reduction process program, G-protein-coupled receptor signaling, and several other metabolic pathways. Of note, many genes (e.g., *Cox6c* and *Cox6a1*) overlap between oxidation-reduction program and OXPHOS program. Quantitative RT-PCR results confirmed that expression of many tumor-promoting program genes was similarly increased by both *Nestin-Cre*-mediated heterozygous loss of *Kmt2d* (**Fig. 3c**) and Atoh1-Cre-mediated heterozygous loss of *Kmt2d* (**Fig. 3d**). Interestingly, expression levels of most of these tumor-promoting genes anti-correlated with *KMT2D* mRNA levels in human MBs, suggesting human MB relevance of our findings (**Fig. 3e**). Consistent with the notion that G-coupled receptor signaling may enhance activities of cellular kinases ^42,43^, IHC analysis showed that heterozygous *Kmt2d* loss increased p-AKT1 and p-ERK levels (**Fig. 3f,g**).

**Figure 3:**
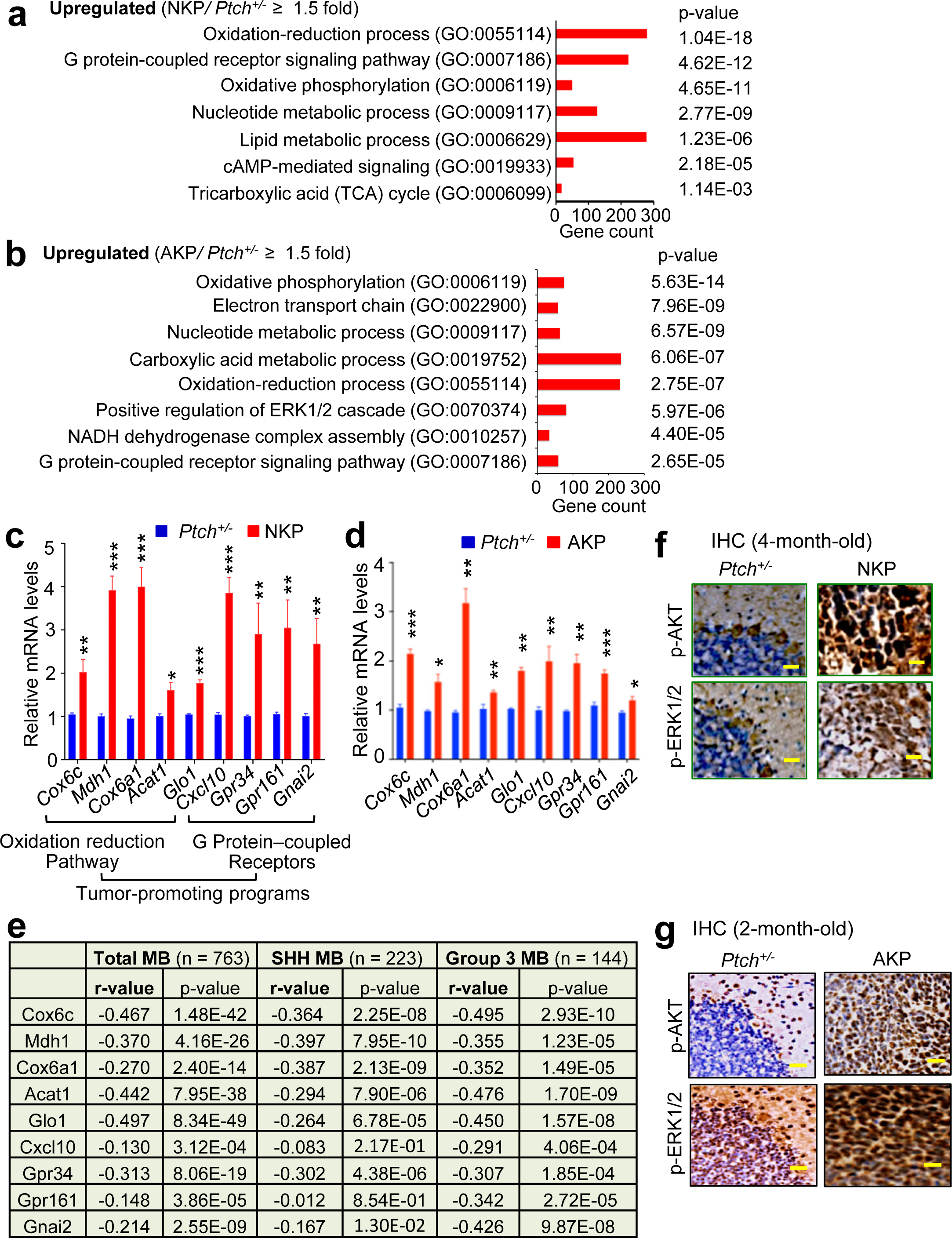
Heterozygous *Kmt2d* loss upregulates tumor-promoting programs, including oxidation-reduction/OXPHOS pathway and G-protein-coupled receptor signaling, for MB genesis. (**<u>a</u> & <u>b</u>**) Ontology analysis of genes that are at least 1.5-fold upregulated in NKP (4-month-old) and AKP (2-month-old) cerebellar tissues over *Ptch^+/−^* cerebellar tissues. Gene ontology analysis was performed using the Panther analysis. (**<u>c</u> & <u>d</u>**) Quantitative RT-PCR analysis of expression levels of oxidation-reduction (including OXPHOS) genes and G protein-coupled receptor genes between *Ptch^+/−^* and NKP cerebellar tissues (**c**) or between *Ptch^+/−^* and AKP cerebellar tissues (**d**). Data are presented as the mean ± SEM (error bars) of at least three independent experiments. *p < 0.05, **p < 0.01, and ***p < 0.001. (**<u>e</u>**) Reverse correlation of KMT2D with oxidation-reduction (including OXPHOS) genes and G protein-coupled receptor genes. Cavalli dataset (n = 763) in the R2 database were analyzed. (**<u>f</u> & <u>g</u>**) Representative IHC images of p-AKT, p-ERK1/2, and KMT2D for *Ptch^+/−^* and NKP cerebellar tissues (**f**) or for *Ptch^+/−^* and AKP cerebellar tissues (**g**). See also **Extended Data Fig. 5**.

In addition, heterozygous *Kmt2d* loss commonly downregulated several gene expression programs (e.g., neurogenesis/neuron development, neuronal differentiation, and synapsis organization) in both NKP and AKP cerebella compared to *Ptch*^+/−^ cerebella (**Extended Data Fig. 5d,e**). Genes in such programs included neuronal differentiation transcription factor genes *Neurod1* and *Neurod2* (**Extended Data Fig. 5f**). Thus, our findings are in line with the notion that tumorigenesis is associated with induced oncogenic programs and dysregulated differentiation ^44,45^.

As aforementioned, heterozygous *Kmt2d* loss significantly upregulated oxidation-reduction process/OXPHOS genes (e.g., Cox6c) and G protein-coupled receptor signaling factor genes (e.g., GPR161 and GPR34). To understand what role these genes play in promoting MB, we determined whether their knockdown affects the proliferation of MB cells. For this, we used the human MB cell line DAOY and also established *Atoh1-Cre Kmt2d^fl/+^ Ptch^+/−^* (called AKP) MB cell line and *Ptch*^+/−^ MB cell line from their respective murine MBs ― these two murine MB cell lines are in the isogenic state. Difference of gene expression between these two murine MB cell lines recapitulated the effect of heterozygous *Kmt2d* loss on gene expression in *Ptch*^+/−^ cerebellum (**Fig. 4a vs 3d**). In addition, consistent with increased expression of OXPHOS genes in the AKP MB cell line, oxygen consumption rate (OCR, a readout of OXPHOS) was increased in the AKP MB cell line compared to the *Ptch*^+/−^ MB cell line (**Fig. 4b**). COX6C is a subunit of the cytochrome C oxidase complex important for electron transport during OXPHOS in the mitochondria; and this protein is involved in promoting tumorigenesis ^46^. In line with this, COX6C knockdown reduced the proliferation of the AKP MB cell line and of human DAOY cells (**Fig. 4c,d**). Moreover, COX6C knockdown reduced OCR of human DAOY cells (**Fig. 4e**). The G protein-coupled receptors GPR161 and GPR34 are capable of promoting tumorigenesis ^42,43,47^. Knockdown of GPR161 and GPR34 decreased cell proliferation (**Fig. 4f,g and Extended Data Fig. 6a,b**), and GPR34 knockdown reduced p-AKT1 and p-ERK levels in AKP MB cell line (**Extended Data Fig. 6c**). These results suggest that upregulation of these genes contributes to the MB-promoting effect of heterozygous *Kmt2d* loss.

**Figure 4:**
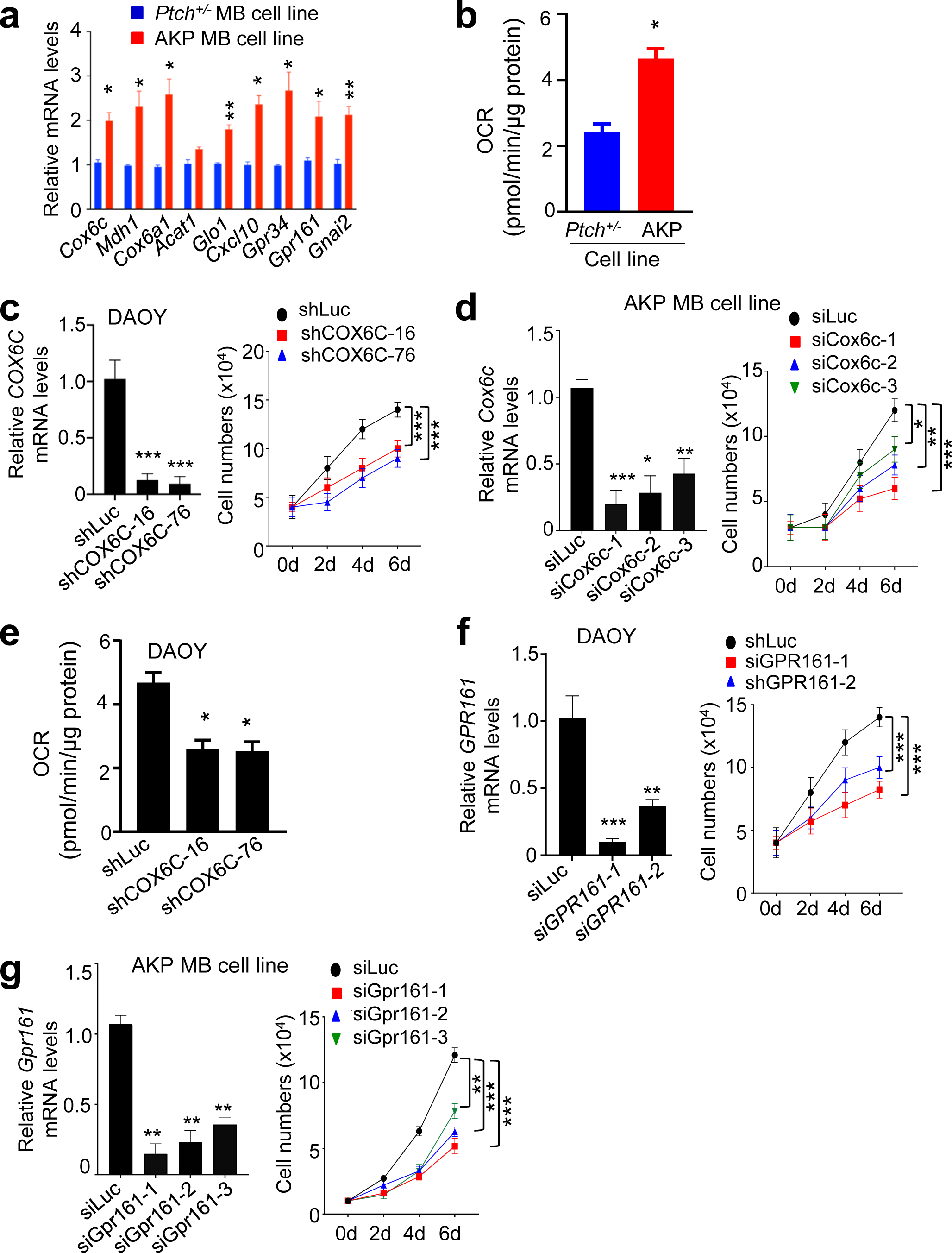
The OXPHOS factor COX6C and the G-protein-coupled receptor GPR161 are required for the proliferation in MB cells. (**<u>a</u>**) Quantitative RT-PCR analysis of expression levels of oxidative phosphorylation-associated genes and G protein-coupled receptor genes in *Ptch^+/−^* and AKP MB cell line. Data represent the mean ± SEM (error bars) of at least three independent experiments or biological replicates. *p < 0.05 and **p < 0.01. (**<u>b</u>**) Comparison of oxygen consumption rate (OCR) between *Ptch^+/−^* and AKP MB cells. (**<u>c</u> & <u>d</u>**) The effect of COX6C knockdown on the proliferation of DAOY (**c**) and AKP MB (**d**) cell line. (**<u>e</u>**) The effect of COX6C knockdown on OCR levels in DAOY cells. (**<u>f</u> & <u>g</u>**) The effect of GPR161 knockdown on the proliferation of DAOY (**f**) and AKP MB (**g**) cell line. Data are presented as the mean ± SEM, *p < 0.05, **p < 0.01 and ***p < 0.001. See also **Extended Data Fig. 6**.

### Heterozygous *Kmt2d* loss downregulates the transcription-repressive tumor suppressor NCOR2 to indirectly upregulate tumor-promoting genes

Although the tumor-promoting programs were upregulated by heterozygous *Kmt2d* loss, KMT2D may not directly repress the tumor-promoting programs because KMT2D generally acts as a transcriptional coactivator. Therefore, we hypothesized that such tumor-promoting programs would be repressed by a transcription-repressive tumor-suppressive factor (s) whose expression is activated by KMT2D. Thus, we searched which transcription-repressive, tumor-suppressive factors were downregulated (≤ 0.5-fold) by heterozygous *Kmt2d* loss. In addition, we incorporated human MB relevance into this search by examining which factors correlate (r ≥ 0.2) with *KMT2D* expression in human MB samples. This search led us to find several candidates (**Fig. 5a,b**). Quantitative RT-PCR confirmed that their expression levels were reduced in NKP and AKP cerebella compared to *Ptch*^+/−^ cerebella (**Fig. 5c,d**). Of these candidates, NCOR2’s expression levels best correlated with *KMT2D* levels in MB samples, including SHH and group 3 MB samples (**Fig. 5b**). In addition, quantitative RT-PCR and IHC staining showed that *NCOR2* expression was highly decreased by heterozygous *Kmt2d* loss (**Fig. 5c–f**).

**Figure 5:**
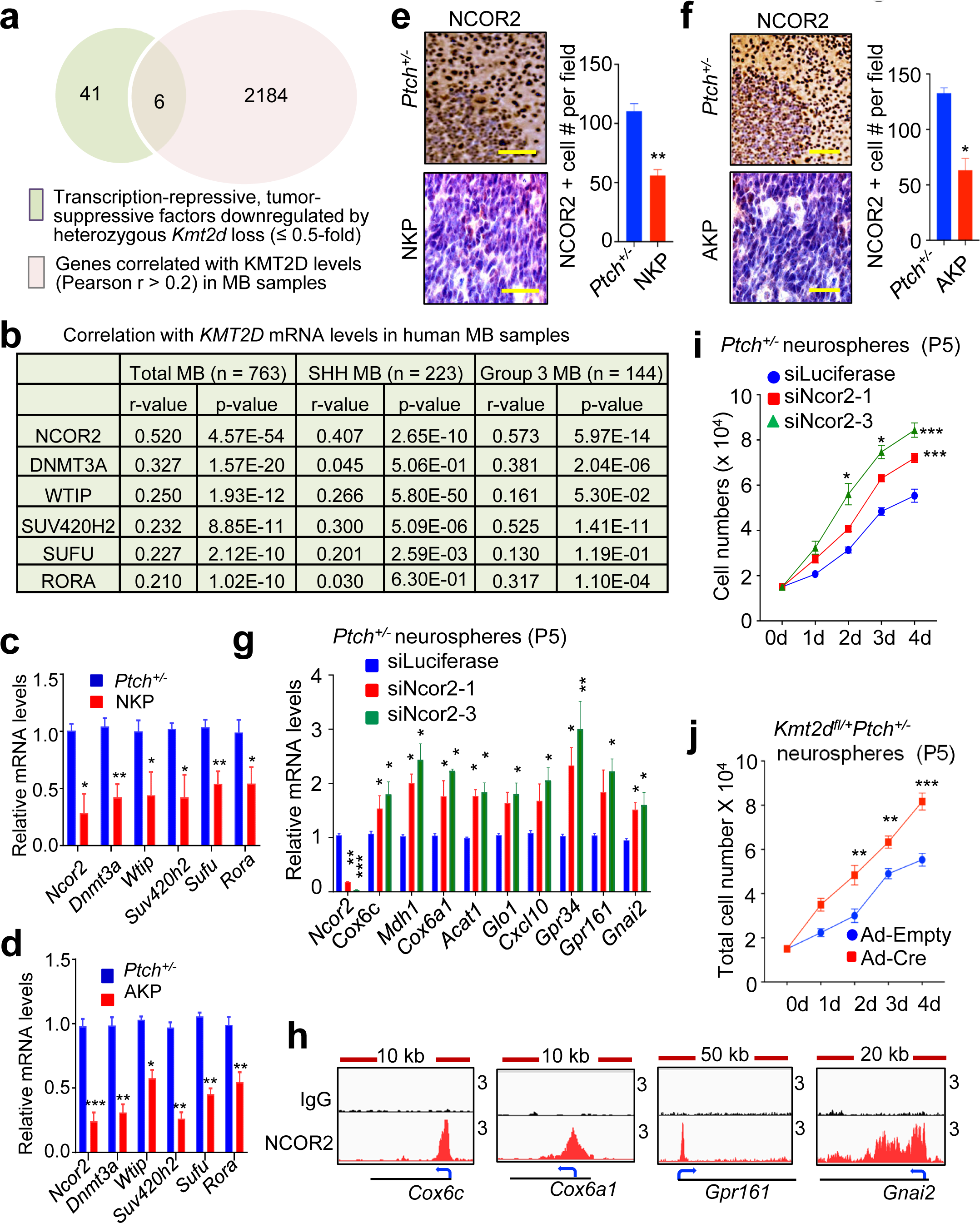
Decreased expression of NOCR2 by heterozygous *Kmt2d* loss upregulates tumor-promoting programs. (**<u>a</u>**) Venn diagram for an overlap between transcription-repressive, tumor-suppressive factor genes downregulated by heterozygous *Kmt2d* loss and genes whose expression correlates with KMT2D levels. Cavalli dataset (n = 763) in the R2 database was used for correlation analysis. (**<u>b</u>**) Significant correlation of *NCOR2* expression with *KMT2D* expression in total human MB, SHH MB, and Group 3 MB samples (Cavalli dataset). (**<u>c</u> & <u>d</u>**) Analysis of *Ncor2* mRNA levels between *Ptch^+/−^* and NKP cerebellar tissues (**c**) or between *Ptch^+/−^* and AKP cerebellar tissues (**d**). (**<u>E</u> & <u>F</u>**) Representative images of NCOR2 IHC staining and quantification of IHC images for *Ptch^+/−^* and NKP cerebellar tissues (**e**) and for *Ptch^+/−^* and AKP cerebellar tissues (**f**). (**<u>g</u>**) The effect of NCOR2 knockdown on gene expresison in *Ptch^+/−^* neurospheres. (**<u>h</u>**) CUT&RUN profiles for IgG (control) and NCOR2 at the *Cox6c*, *Cox6a1*, *GPR161*, and *Gnai2* genes in the *Ptch^+/−^* cerebellar tissues. (**<u>i</u>**) The effect of NCOR2 knockdown on the proliferation of *Ptch^+/−^* neurospheres. (**<u>j</u>**) The effect of heterozygous *Kmt2d* loss on *Kmt2d^fl/+^ Ptch^+/−^* neurospheres. Data represent the mean ± SEM, *p < 0.05, **p < 0.01 and ***p < 0.001.

NCOR2 (nuclear receptor corepressor 2; also called SMRT) is a transcriptional corepressor that regulates chromatin and gene expression ^48^. NCOR2 can act as an anti-tumorigenic factor in several types of cancer. For instance, *NCOR2* is downregulated in multiple myeloma, and its loss is linked with the neoplastic transformation of non-Hodgkin’s lymphoma ^49,50^. We have shown that NCOR2 represses expression of tumor-promoting factors in lung cancer ^51^. *NCOR2* harbors some nonsense mutations in MBs ^22^, implicating its tumor-suppressive function. Interestingly, NCOR2 knockdown in P4‒P6 *Ptch*^+/−^ neurospheres upregulated genes for G-protein-coupled receptor signalling and oxidation-reduction process program (**Fig. 5g**). NOCR2 occupied such tumor-promoting genes, indicating that NCOR2 directly represses them (**Fig. 5h**). NCOR2 knockdown increased the proliferation of *Ptch*^+/−^ neurospheres, and this effect of NCOR2 knockdown was comparable to that of heterozygous *Kmt2d* loss in *Ptch*^+/−^ neurospheres (**Fig. 5i,j**). These results indicate that NCOR2 is a major effector downstream of KMT2D’s tumor-suppressive function.

### The onco-protein MycN, which is upregulated by heterozygous *Ptch* loss, activates tumor-promoting genes

Considering that heterozygous *Kmt2d* loss and heterozygous *Ptch* loss cooperate for MB genesis, we also investigated how heterozygous *Ptch* loss contributes to this cooperation. As it has been known that heterozygous *Ptch* loss can upregulate oncogenic factors in MB ^52^, we rationalized that an oncogenic factor(s) upregulated by heterozygous *Ptch* loss would play a role in this cooperation by activating tumor-promoting genes. We searched for SHH-PTCH’s downstream onco-proteins that are upregulated in NKP cerebella and AKP cerebella. Interestingly, we found that *MycN* (alias *NMyc*) was highly upregulated in NKP and AKP cerebella (**Fig. 6a,b**). Consistent with this, IHC levels of MycN were increased in NKP and AKP cerebella (**Fig. 6c,d**). *MycN* is often amplified and overexpressed in MBs ^53,54^, is a major transcription activator downstream of SHH-PTCH pathway, and is a potent oncogene for MB genesis ^55,56^. In line with *MycN* upregulation, *E2f1* and *E2f2,* both of which are MYCN’s target genes ^52^, were also upregulated in NKP and AKP cerebella (**Fig. 6a,b**). E2F1 and E2F2 positively regulate cell cycle. E2F1 overexpression in brain induced a brain tumor ^57^.

**Figure 6:**
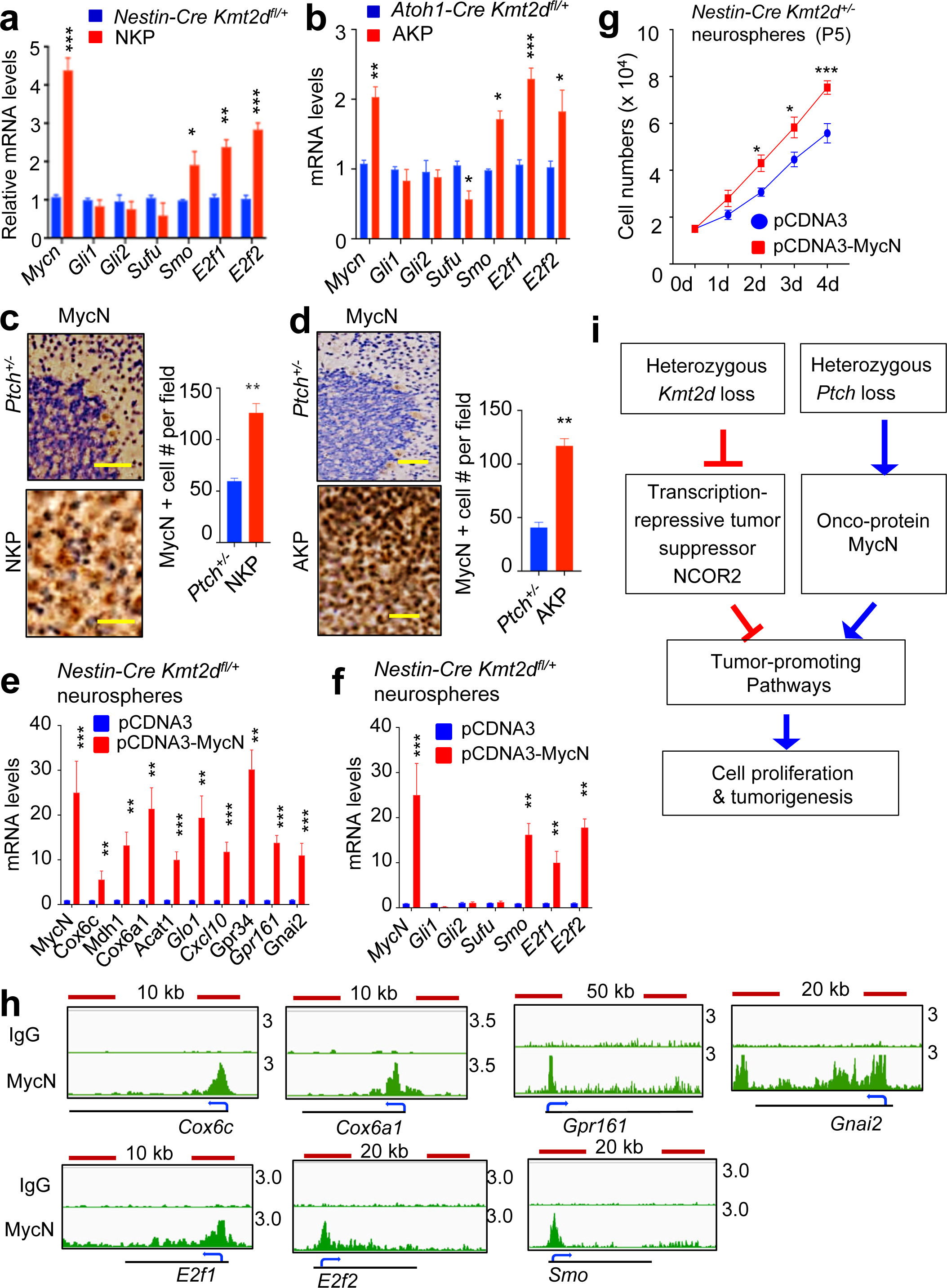
Upregulation of MycN by heterozygous *Ptch* loss directly activates tumor-promoting genes. (**<u>a</u> & <u>b</u>**) Comparison of *Mycn*, *Gli1*, *Gli2*, *Sufu*, *Smo*, *E2f1*, and *E2f2* mRNA levels between *Nestin-Cre Kmt2d^fl/+^* and NKP cerebella (4-month-old) (**a**) or between *Atoh1-Cre Kmt2d^fl/+^* and AKP cerebella (2-month-old) (**b**). (**<u>c</u> & <u>d</u>**) Comparison of IHC images and quantification of MycN between *Ptch^+/−^* and NKP cerebellar tissues (**c**) or between *Ptch^+/−^* and AKP cerebella (**d**). (**<u>e</u> & <u>f</u>**) The effect of ectopic expression of MycN on tumor-promoting genes in *Nestin-Cre Kmt2d^fl/+^* neurospheres. (**<u>g</u>**) The effect of MycN overexpression on the proliferation assays of *Nestin-Cre Kmt2d^fl/+^* neurospheres. Data are presented as the mean ± SEM. *p < 0.05, **p < 0.01 and ***p < 0.001. (**<u>h</u>**) CUT&RUN analysis for IgG (control) and MycN at the *Cox6c*, *Cox6a1*, *GPR161*, *Gnai2, E2f1, E2f2,* and *Smo* genes in the *Ptch^+/−^* mouse cerebellum. (**I**) The proposed mechanistic pathways showing MB formation by heterozygous *Kmt2d* loss and heterozygous *Ptch* loss.

As it has been unknown whether MycN regulates the aforementioned tumor-promoting genes in MB, we assessed the effect of MycN overexpression on the tumor-promoting genes. For this, we ectopically expressed *MycN* in P4‒P6 cerebellar cells (e.g., *Nestin-Cre Kmt2d^fl/+^* neurospheres). *MycN* overexpression increased the expression of the tumor-promoting genes in *Nestin-Cre Kmt2d^fl/+^* neurospheres and upregulated the proliferation of these neurospheres (**Fig. 6e–g**). Our ChIP-seq results showed that MycN occupied the tumor-promoting genes, indicating that MycN directly activates them (**Fig. 6h**). Taken together, these findings point out that *MycN* upregulation by heterozygous *Ptch* loss cooperates with *NCOR2* downregulation by heterozygous *Kmt2d* loss to cooperatively active tumor-promoting genes for MB genesis (**Fig. 6i**).

### Heterozygous *Kmt2d* loss impairs super-enhancers/enhancers and H3K4me3 signatures at both the genome-wide level and tumor suppressor genes

Others and we have shown that KMT2D is able to *in vitro* catalyze mono-, di- or trimethylation at H3K4 ^58–60^ and that KMT2D generates the enhancer mark H3K4me1 in cells ^61–63^ and also helps produce another enhancer mark acetylated H3K27 (H3K27ac) ^64,65^ to activate gene expression in cells. We have also shown that KMT2D activates gene expression by engendering H3K4me3 at gene promoters ^58,59,66^ and that homozygous *Kmt2d* loss downregulates H3K4me1, H3K27ac and H3K4me3 peaks to decrease gene expression ^9^. For these reasons, we examined whether heterozygous *Kmt2d* loss alters H3K4me1, H3K27ac, and H3K4me3 in *Ptch*^+/−^-driven MB. Unexpectedly, IF analysis showed that heterozygous *Kmt2d* loss highly decreased nuclear levels of H3K4me1, H3K27ac, and H3K4me3 in NKP and AKP cerebella compared to *Ptch*^+/−^ cerebella (**Fig. 7a,b and Extended Data Fig. 7a,b**).

**Figure 7:**
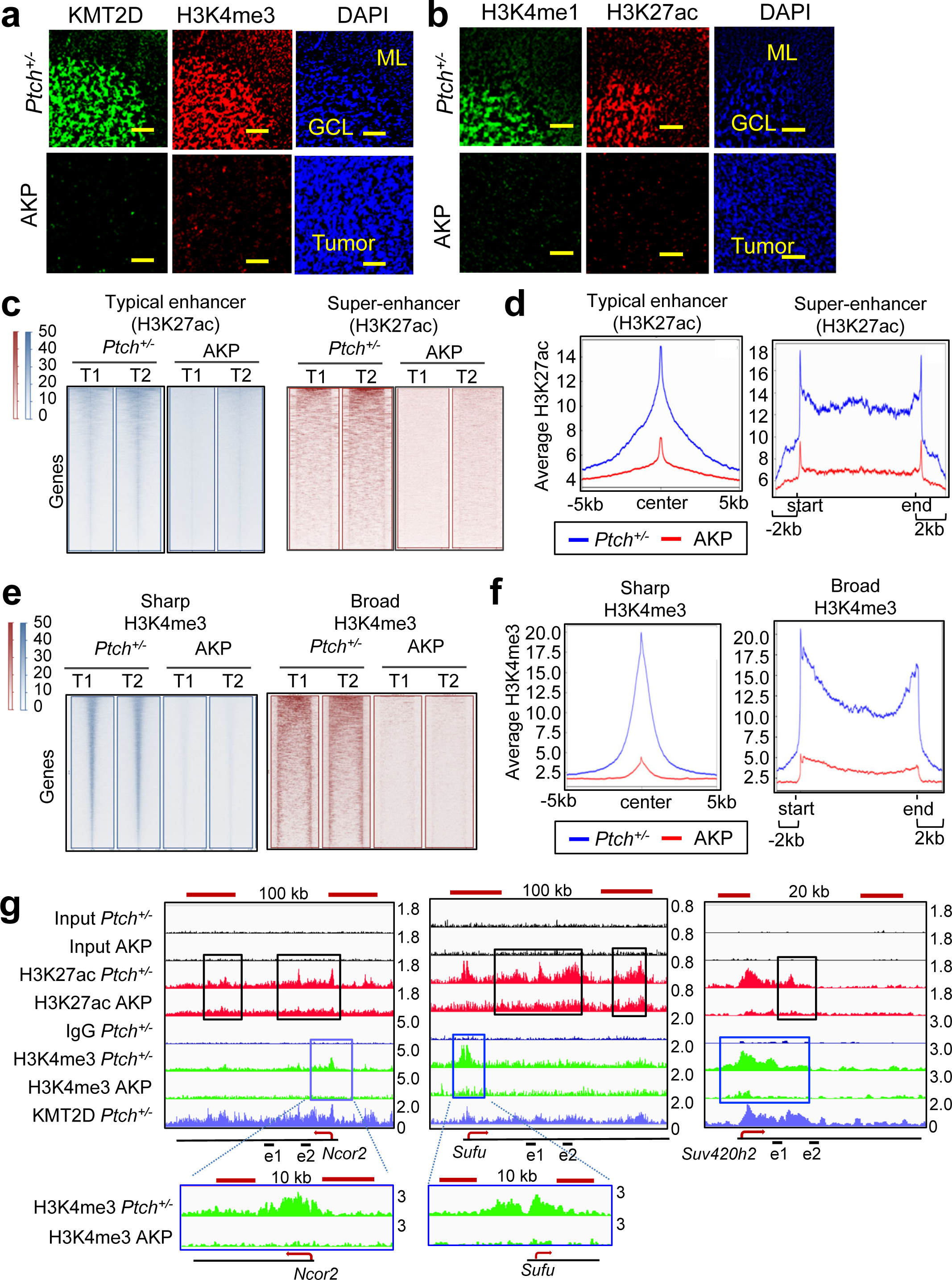
Heterozygous *Kmt2d* loss downregulates enhancers and H3K4me3 signatures at tumor suppressor genes. (**<u>a</u> & <u>b</u>**) Immunofluorescence (IF) analysis for KMT2D, H3K4me3, H3K4me1, and H3K27ac in *Ptch^+/−^* and *Atoh1-Cre Kmt2d^fl/+^ Ptch^+/−^*(AKP) cerebella. (**<u>c</u> & <u>d</u>**) Heat maps of ChIP-seq read (RPKM) for H3K27ac signals (**c**) and average H3K27ac intensity curves (**d**) at typical enhancer and super-enhancer regions in *Ptch^+/−^* and AKP cerebella (2-month-old). (**<u>e</u> & <u>f</u>**) Heat maps of CUT&RUN read (RPKM) for H3K4me3 signals (**c**) and average H3K4me3 intensity curves (**d**) at sharp and broad H3K4me3 regions in *Ptch^+/−^* and AKP cerebella (2-month-old). The intensity scale of the color is from 0 to 50. (**<u>g</u>**) ChIP-seq profiles for H3K27ac (red) and CUT&RUN peaks for H3K4me3 (green), KMT2D (purple), and IgG (blue) at the *Ncor2*, *Sufu*, and *Suv420h2* genes in *Ptch^+/−^* and AKP cerebellar tissues (2-month-old). Signals in the tracks were generated by subtracting input or IgG signals from their original signals. See also **Extended Data Fig. 7**.

Enhancers are a key type of gene-activating signature that spatiotemporally activates gene expression in distant locations and are marked by H3K4me1 and H3K27ac ^67^. Super-enhancers are clusters of enhancers, which are much more extensive than typical enhancers (a median size of 0.7–1.3 kb). Super-enhancers highly upregulate gene expression and can be linked to activation of cell identity genes and disease-relevant genes ^68–70^. Broad H3K4me3 (also called H3K4me3 breadth) is another gene-activating epigenomic signature that occupies many critical tumor suppressor genes and cell identity genes and is associated with highly active genes ^71,72^. Broad H3K4me3 peaks are characterized by a skewed distribution spanning at least −500 bp to +3,500 bp, which includes the promoters and transcription start sites. They are different from sharp H3K4me3 peaks, which have narrow and high features.

To determine the effect of heterozygous *Kmt2d* loss on genome-wide landscapes of enhancers and H3K4me3 signatures, we performed ChIP-seq using *Ptch*^+/−^ cerebella and AKP cerebella bearing MB. Our enhancer analysis on the basis of H3K27ac’s ChIP-seq signals showed that heterozygous *Kmt2d* loss reduced H3K27ac signals in super-enhancers (top 1000 enhancers) and typical enhancers (**Fig. 7c,d**). In addition, analysis on the basis of H3K4me3’s CUT&RUN signals showed that heterozygous *Kmt2d* loss reduced H3K4me3 signals in broad and sharp H3K4me3 signatures (**Fig. 7e,f**). These results indicate that heterozygous *Kmt2d* loss negatively impact enhancers and H3K4me3 signatures.

To assess the effect of heterozygous *Kmt2d* loss on enhancers and H3K4me3 peaks at individual genes, we analyzed ChIP-seq profiles of H3K27ac and H3K4me3 at the transcription-repressive and tumor-suppressive factor genes (e.g., *Ncor2, Sufu*, and *Suv420h2*) in *Ptch*^+/−^ cerebella and AKP cerebella bearing MB. Our ChIP-seq results showed that *Ncor2* and *Sufu* were occupied by or were close to large domains that contain a cluster of H3K27ac peaks. As these domains range from 50 kb to 100 kb, they would be considered super-enhancers (See black-outlined boxes in **Fig. 7g**) ^68,69^. *Suv420h2* had a typical enhancer (**Fig. 7g**). In addition, the promoter regions of *Ncor2, Sufu*, and *Suv420h2* were decorated by H3K4me3 peaks (See blue-outlined boxes in **Fig. 7g**). Interestingly, heterozygous *Kmt2d* loss decreased enhancer peaks (H3K27ac peaks) and broad H3K4me3 peaks at *Ncor2, Sufu*, and *Suv420h2* (**Fig. 7g**). Importantly, KMT2D occupied enhancers and H3K4me3 peaks in *Ncor2, Sufu*, and *Suv420h2* genes in *Ptch*^+/−^ cerebella (**Fig. 7g**). In enhancers, enhancer RNAs (eRNAs) are transcribed by RNA Polymerase II in a bidirectional manner, and their levels represent enhancer activities ^73^. We examined the effect of heterozygous *Kmt2d* loss on eRNA expression from the enhancers of *Ncor2, Sufu*, and *Suv420h2*. Because these enhancers are located in the gene body, we measured anti-sense eRNAs to distinguish them from mRNAs. Our quantitative RT-PCR results showed that heterozygous *Kmt2d* loss reduced eRNA levels in the enhancers in *Ncor2, Sufu*, and *Suv420h2, suggesting that* heterozygous *Kmt2d* loss decreased enhancer activity. (**Extended Data Fig. 7c**). These results indicate that KMT2D directly and positively regulates super-enhancers/typical enhancers and H3K4me3 peaks of *Ncor2, Sufu*, and *Suv420h2* genes and that heterozygous *Kmt2d* loss downregulates these genes by weakening these epigenomic signatures.

### Pharmacological inhibition impedes the tumorigenic growth of MB cells bearing heterozygous *Kmt2d* loss

To identify a pharmacological strategy for inhibition of the tumorigenic growth of MB cells bearing heterozygous *Kmt2d* loss, we searched small molecule inhibitors on the basis of our aforementioned mechanistic findings. As described above, heterozygous *Kmt2d* loss decreased enhancer signals (H3K4me1 and H3K27ac) and H3K4me3 levels and upregulated OXPHOS program and p-AKT and p-ERK levels. Thus, we sought to pharmacologically inhibit H3K4me1 reduction, H3K27ac reduction, H3K4me3 reduction, OXPHOS program, and p-AKT and p-ERK in MB cells bearing heterozygous *Kmt2d* loss. To inhibit H3K4me1 reduction (i.e., to increase H3K4me1 levels), we used an inhibitor (ORY-2001) of the H3K4me1/H3K4me2 demethylase LSD1, which is critical for decommissioning enhancers ^74^. To increase H3K27ac, we used histone deacetylase (HDAC) inhibitors, such as SAHA (also known as Vorinostat) and Romidepsin, both of which have been approved for cancer treatment by the U.S. Food and Drug Administration (FDA). To increase H3K4me3 levels, we used an inhibitor (KDM5-C70) of the KDM5 family of H3K4me3/me2 demethylases. To inhibit OXPHOS, we used the FDA-approved OXPHOS/diabetes inhibitor Metformin. To inhibit p-AKT and p-ERK, we used TIC10 (a dual inhibitor of AKT/ERK, also called ONC201), an ERK inhibitor (SCH772984), and an AKT inhibitor (Capivasertib/AZD5363 in clinical trials of Phase III).

To test the effects of these inhibitors on proliferation of MB cells bearing heterozygous *Kmt2d* loss, we determined which inhibitor preferentially impedes the proliferation of our AKP MB cell line compared to that of our *Ptch*^+/−^ MB cell line. The *Ptch*^+/−^ and AKP MB cell line in the isogenic state had similar cell proliferation rate (**Extended Data Fig. 8a**). ORY-2001 (and to a lesser extent Metformin, TIC10, SCH772984, and Capivasertib) preferentially inhibited the proliferation of the AKP MB cell line compared to that of the *Ptch^+/−^* MB cell line (**Fig. 8a–c, Extended Data Fig. 8b,c**). In contrast, SAHA, Romidepsin, and KDM5-C70 did not demonstrate obvious selective inhibition of the proliferation of the AKP MB cell line compared to that of *Ptch^+/−^* MB cell line (**Extended Data Fig. 8d–f**).

**Figure 8:**
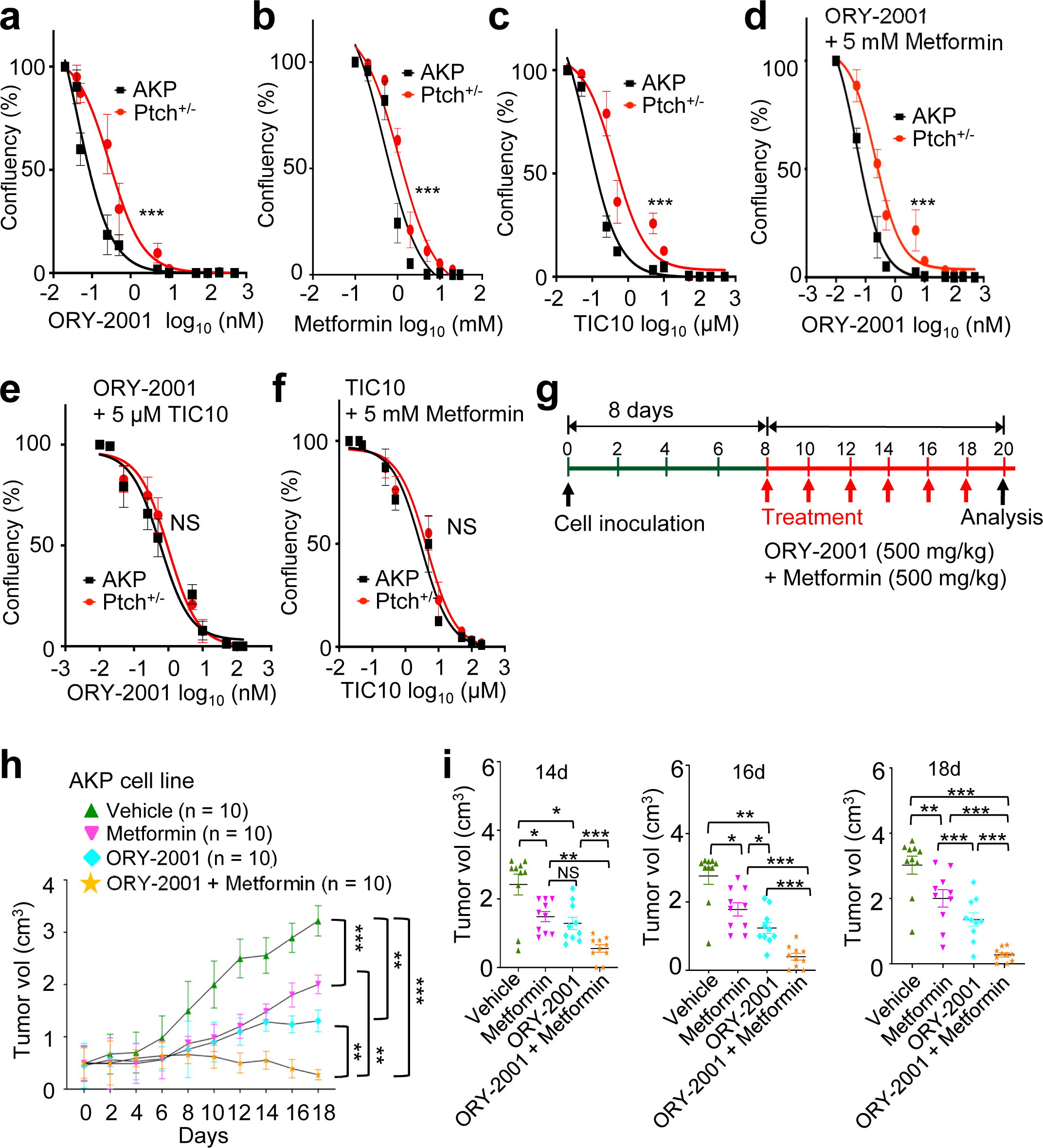
Combinatory treatment of ORY-2001 and Metformin eradicates tumor formation of MB cells with heterozygous *Kmt2d* loss. (**<u>a</u>‒<u>f</u>**) The effects of inhibitors ORY-2001, Metformin, and TIC10 on the proliferation of *Ptch^+/−^* and AKP cell lines. Cells were treated with various concentrations of ORY-2001 (**a**), Metformin (**b**), TIC10 (**c**), ORY-2001 + 5 mM Metformin (**d**), ORY-2001 + 5 µM TIC10 (**e**), and TIC10 + 5 mM Metformin (**f**) for 72h. (**<u>g</u>‒<u>i</u>**) The effect of ORY-2001, Metformin, and their combination (ORY-2001 + Metformin) on tumorigenic growth of AKP MB cells in a mouse subcutaneous xenograft model. The schedule of treatment of mice is shown (**g**). The sizes of xenograft tumors were measured (**h**). The individual tumor volumes for 14d, 16d, and 18d are shown separately (**i**). Data are presented as the mean ± SEM (error bars) of at least three independent experiments or biological replicates. *p < 0.05, **p < 0.01, and ***p<0.001. See also **Extended Data Fig. 8**.

Combinatory drug therapies are often used for more effective cancer treatment. Since three types of inhibitors (LSD1 inhibitor ORY-2001, Metformin, and AKT/ERK inhibitors) preferentially impeded the proliferation of the AKP MB cell line, we examined whether combination of three inhibitors (ORY-200, Metformin, and the dual AKT/ERK inhibitor TIC10) synergistically inhibits the proliferation of the AKP MB cell line than do single inhibitors. Only the combination of ORY-2001 and Metformin preferentially inhibited the proliferation of the AKP MB cell line compared to that of the *Ptch^+/−^* MB cell line (**Fig. 8d–f**). We also examined the combinatory effect of ORY-2001 and Metformin on tumorigenic growth of the AKP MB cells in a subcutaneous mouse model. This combination almost completely inhibited the tumorigenic growth of AKP MB cells whereas individual drugs weakly did (**Fig. 8g–i and Extended Data Fig. 8g**). ORY-2001 and Metformin did not have a significant effect on mouse body weight, suggesting no major toxicity of this combinatory treatment (**Extended Data Fig. 8h**). These results indicate that the tumorigenic growth of MB cells bearing heterozygous *Kmt2d* loss can be effectively eradicated by pharmacological combinatory inhibition of LSD1 and OXPHOS.

## DISCUSSION

In the present study, our results revealed that heterozygous *Kmt2d* loss in the brain or the cerebellum-containing local region highly promoted *Ptch^+/−^*-driven MB genesis in mice, indicating a critical role for heterozygous loss of the potent MB suppressor KMT2D in MB genesis. Considering that most of the *KMT2D* mutations in human MB are heterozygous, this finding would be clinically relevant to human MB. Mechanistically, downregulation of the transcription-repressive tumor suppressor gene *NCOR2* by heterozygous *Kmt2d* loss cooperates with upregulation of the oncogene *MycN* by heterozygous *Ptch* loss to increase the expression of tumor-promoting genes (e.g., the OXPHOS gene *Cox6c* and G protein-coupled receptor signaling factor genes *GPR161 and GPR34*). Notably, heterozygous *Kmt2d* loss drastically downregulated super-enhancers/enhancers and H3K4me3 signature at tumor suppressor genes (e.g., *Ncor2*) to reduce their expression. Our findings uncover an original MB-promoting epigenetic mechanism in which heterozygous loss of a tumor-suppressive epigenetic modifier downregulates other tumor suppressor genes via downregulation of gene-activating epigenomic signatures and thereby upregulates tumor-promoting programs to accelerate MB genesis.

*KMT2D* and *PTCH* are two of the most frequently mutated tumor suppressor genes in MB. KMT2D is an epigenetic modifier that catalyzes histone lysine methylation. PTCH is a tumor-suppressor in SHH signaling. Our results showed that KMT2D directly and positively regulated *Ncor2* expression via activation of enhancer and H3K4me3 signature. It has been known that PTCH downregulates the oncoprotein MycN’s levels ^56^. Our results indicate that NCOR2 acts as a tumor suppressor by occupying and repressing tumor-promoting genes, whereas MycN occupies and positively regulates similar tumor-promoting genes. These findings highlight an unexpected link between the epigenetic tumor suppressor KMT2D and the SHH signaling suppressor PTCH as heterozygous *Kmt2d* loss and heterozygous *Ptch* loss commonly activate a similar set of tumor-promoting genes via their downstream onco-factors. Thus, our findings provide molecular insights into how an aberrant epigenetic modifier is connected with altered SHH signaling pathway to cooperate for MB genesis.

Super-enhancers are associated with cancer ^68,70,75^. This gene-activating signature often upregulates oncogenes in cancer cells ^76^. Therefore, it has been suggested that oncogenes can be inactivated by disruption of super-enhancers by BET-bromodomain inhibitors (e.g., JQ1) ^68^. In contrast, we previously showed that tumor suppressor genes could be inactivated by the impairment of tumor-suppressive super-enhancers by homozygous *Kmt2d* loss ^69^. In line with this, the present study showed that even heterozygous *Kmt2d* loss strongly downregulated epigenomic signals of super-enhancers at tumor suppressor genes. Therefore, although oncogenic super-enhancers would promote cellular transformation during tumorigenesis, certain KMT2D-activated super-enhancers play an important role in MB suppression and would be downregulated by heterozygous *Kmt2d* loss. This tumor-suppressive view of super-enhancers should be carefully considered when a treatment strategy using super-enhancer inhibition would be explored.

Our results indicate that G-protein-coupled receptor signaling and oxidation-reduction process/oxidative phosphorylation program are among tumor-promoting programs induced by heterozygous *Kmt2d* loss. Notably, these programs are largely distinct from those upregulated by homozygous *Kmt2d* loss (e.g., Ras activators and Notch pathway) ^9^. In addition, our results showed that heterozygous *Kmt2d* loss downregulated transcription-repressive tumor suppressor genes (e.g., *Ncor2* and *Suv420h2*) and that NCOR2 decreased expression of several tumor-promoting genes. This effector (i.e., NCOR2) downstream of heterozygous *Kmt2d* loss is somewhat different from major effectors (SIRT1, BCL6 and DNMT3A) downstream of homozygous *Kmt2d* loss ^9^. Thus, MB-promoting mechanisms regulated by heterozygous *Kmt2d* loss appear to be distinct from those regulated by homozygous *Kmt2d* loss and represent a previously unappreciated molecular pathogenesis for MB.

We generated two new MB mouse models (*Nestin-Cre Kmt2d*^fl/+^ *Ptch*^+/−^ mice and *Atoh1-Cre Kmt2d*^fl/+^ *Ptch*^+/−^ mice). Our results using both models commonly showed that heterozygous *Kmt2d* loss remarkably accelerated MB genesis driven by heterozygous *Ptch* loss. Moreover, both *Atoh1*-*Cre*-mediated and *Nestin-Cre*-mediated heterozygous loss of *Kmt2d* promoted *Ptch^+/−^-*driven MB genesis by upregulating similar tumor-promoting programs. As *Atoh1*-*Cre* removes a floxed gene largely in cerebellar GC lineage while *Nestin-Cre* deletes it throughout the brain, these results indicate that MB-promoting effect of heterozygous *Kmt2d* loss rather autonomously occurs in cerebellar GC lineage but is unlikely impacted by heterozygous *Kmt2d* loss in other parts of the brain.

The *KMT2D* gene is mutated and deleted in many other types of cancer, such as bladder, colorectal, oesophageal-gastric, head-neck, and lung cancer as well as melanoma and lymphoma. Notably, heterozygous mutations of *KMT2D* are prevalent in these types of cancer. However, whether and how heterozygous *Kmt2d* loss promotes these cancers are largely unknown. In this regard, our findings regarding the molecular basis underlying the effect of heterozygous *Kmt2d* loss on MB genesis would have significant implications for future studies about how heterozygous *KMT2D* loss orchestrates the genesis of other types of cancer and cooperates with other genetic and epigenetic alterations.

Current standard treatment of MB, including surgery, chemotherapy, and radiation therapy, frequently causes severe neurological deficits and needs to be improved ^77^. In addition, most MBs are negative for PD-L1 and have minimal T cell infiltration. Thus, although anti-PD-L1/anti-PD-1 are in clinical trials for MB patients, patients may have a limited benefit from these immune checkpoint inhibitors [Reviewed in ^78^]. Therefore, there is a great need for a new mechanistic understanding that may eventually help design an effective mechanism-based approach for MB treatment. Our therapeutic strategy using combinatory pharmacological inhibition of LSD1 and OXPHOS of MBs bearing heterozygous *Kmt2d* loss can be relevant to a treatment approach for KMT2D-deficient MBs. Interestingly, the OXPHOS program as essential mitochondrial bioenergetics is indispensable for many types of cancer and can be considered a potential target for cancer therapy ^79^. LSD1 negatively regulates enhancers via H3K4 demethylation, and some LSD1 inhibitors are in clinical trials for the treatment of other types of cancer ^80^. In this regard, as heterozygous *KMT2D* loss appears to be widespread in other types of cancer, this strategy may be applicable to therapeutic approaches for other types of KMT2D-deficient cancer.

## ACKNOWLEDGMENTS

We are thankful to Dr. Z. Han for technical assistance and to the Small Animal Imaging Facility (supported by the NIH/NCI under Core grant award number P30CA16672) at The University of Texas MD Anderson Cancer Center for technical advice. We also thank the IDDRC neuropathology core (Core grant award number U54HD083092) and RNA In Situ Hybridization core at The Baylor College of Medicine for histology and image scan services. L.R. and C.Z. were supported by the CPRIT Research Training Award CPRIT Training Program (RP210028). C.Z. were also supported by the 1R25CA240137-01A1 UPWARDS Training Program (Underrepresented Minorities Working Towards Research Diversity in Science). This work was supported in part by grants to M.G.L. from the NIH (R01CA 262324) and the MD Anderson and by an internal philanthropy funds to S.S.D. from the MD Anderson.

## AUTHOR CONTRIBUTIONS

S.S.D. planned and performed experiments, analyzed data, prepared figures, and wrote the manuscript. C.B., A.Z., L.R., S.K., C.Z., J.A., J-I. A., V.R., and K.C. performed experiments and/or analyzed and interpreted data. M.G.L. conceived the study, designed and directed the study, analyzed data, and wrote the manuscript.

## DECLARATION OF INTERESTS

S.S.D and M.G.L. are inventors of a provisional patent application (pending) entitled Methods and Compositions for the Treatment of KMT2D-deficient Cancers (63/590,682), which has been submitted to the US Patent and Trade Office. The other authors declare no competing interests.

---

**Supplementary Table 1.**
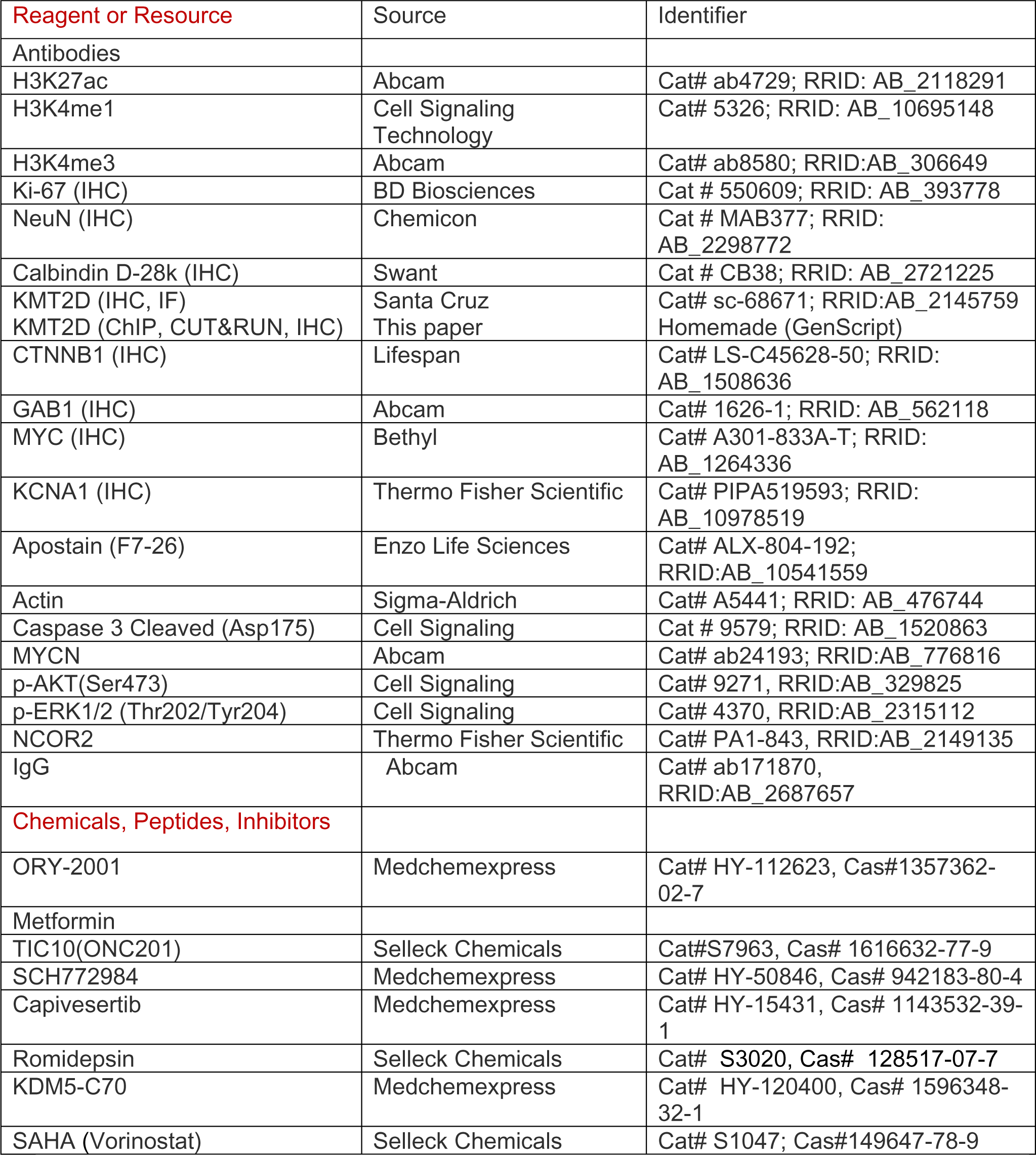
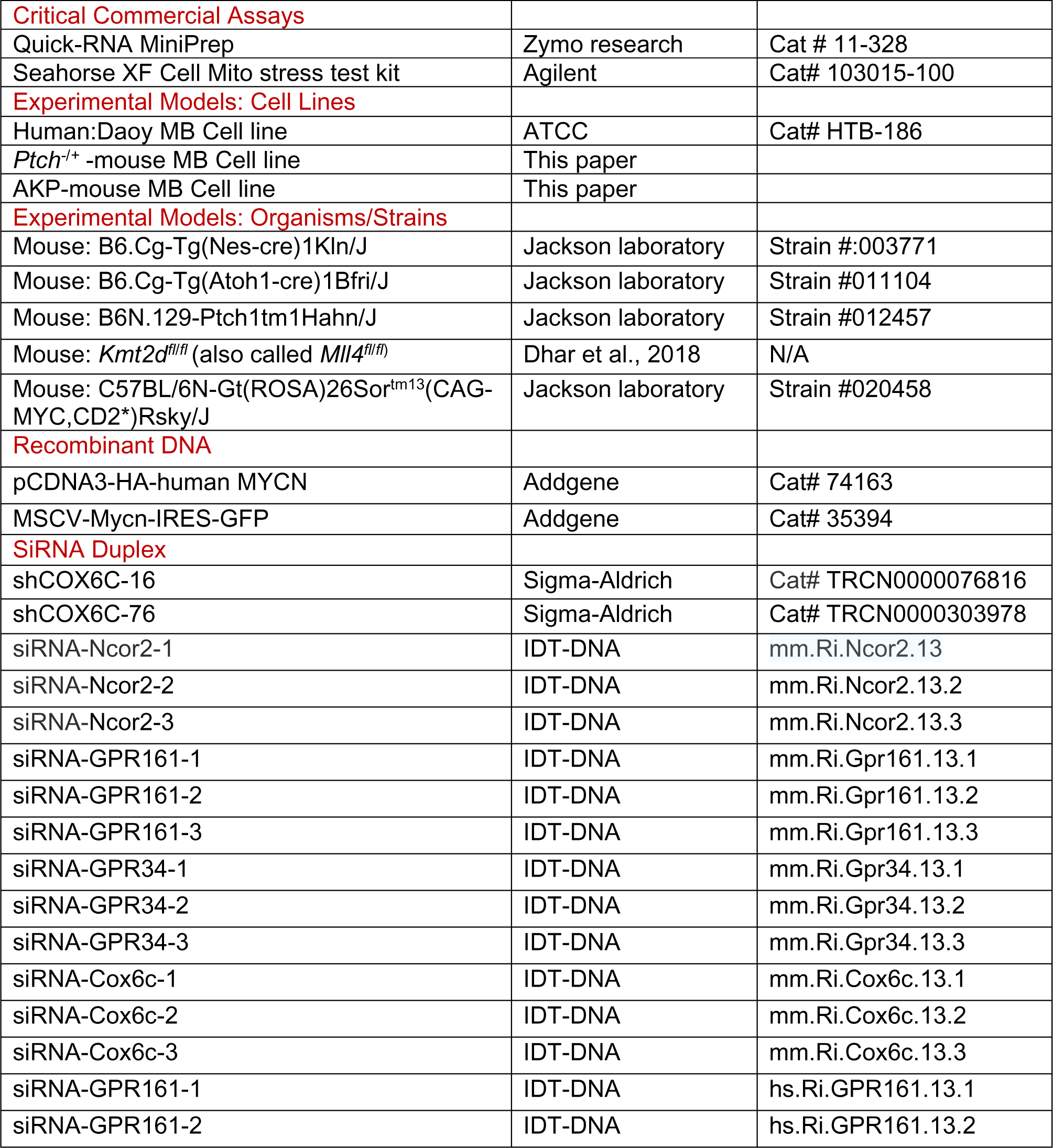
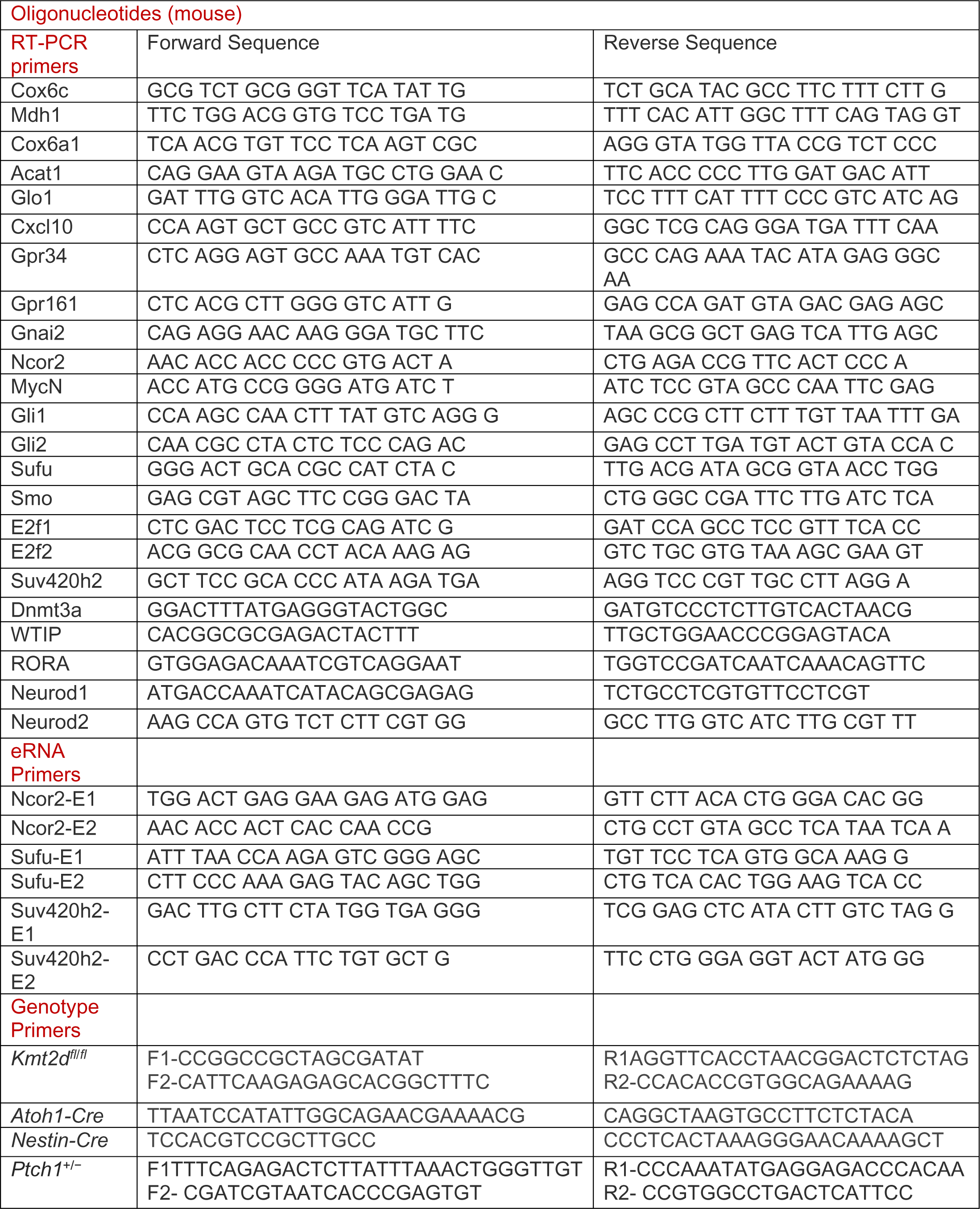
Key resources.

## METHODS

### Mouse Models

Genetically engineered mice used here were *Nestin* (*Nes)-Cre* (003771, JAX), *Ptch*^+/−^ (003081, JAX), *Atoh1-Cre* (011104, JAX) and *Nes-Cre Kmt2d^fl^*^/+^ mice ^9^. *Nes-Cre Kmt2d*^fl/+^ males were bred to *Ptch*^+/−^ female mice to obtain *Nes-Cre Kmt2d^fl/+^ Ptch^+/−^* (NKP) mice. Similarly, *Atoh1-Cre* male mice were bred to generate *Atoh1-Cre Kmt2d^fl/+^ Ptch^+/−^* (AKP) or Atoh1-Cre Kmt2d^fl/+^mice. All mice were maintained on a mixed C57BL/6 × 129Sv background. The generation of *Nestin-Cre Kmt2d^fl/fl^* (*Mll4* BSKO) or *Nestin-Cre Kmt2d^fl/+^* mice was previously described ^9^. The mice used in the study were kept in a laboratory facility accredited by the American Association of Laboratory Animal Care. They were treated in compliance with the National Institutes of Health guidelines.

### Mouse Study Approval

The Institutional Animal Care and Use Committee (IACUC) of The University of Texas MD Anderson Cancer Center approved the use and study of all the animals involved. Survival curves and median survival were plotted and calculated using GraphPad Prism 9.

### Magnetic Resonance Imaging (MRI)

The BioSpec 70/30 MRI machine from Bruker Corp (Billerica, MA) was utilized to monitor tumor growth. MRI scans were conducted on mice aged 1 to 12 months old. The volumes of cerebellar tumors (2.5 mm³ to 15 mm³) in mice were determined by analyzing the MRI data.

### Immunohistochemical and Immunofluorescence Analysis

Mice aged between 2 and 4 months old were perfused with a 4% paraformaldehyde (PFA) solution to fix the tissues. Subsequently, serial coronal sections with a thickness of 40 μm were cut using a cryostat, and these sections were collected as free-floating samples. For the histological analysis of paraffin sections, which were cut to a thickness of 8 μm, a standard hematoxylin/eosin staining procedure was performed. Immunohistochemistry (IHC) or Immunofluorescence (IF) analysis was carried out. The paraffin sections underwent antigen retrieval using an antigen retrieval solution, followed by blocking in a solution containing 3% BSA and 0.1% Triton X-100 for 1 hour. The primary antibodies used are specified in the key resources table. Finally, staining was quantified using the Aperio Nuclear Algorithm software after scanning the images using Aperio Imagescope (Leica Biosystems).

### RNA-seq and Quantitative RT-PCR

Total RNA was isolated from cerebellar tissues using Trizol reagent (ThermoFisher Scientific, Waltham, MA). RNA samples were sequenced by using the Illumina HiSeq 2000. Raw FASTQ files were processed using the nf-core/rnaseq pipeline (v3.7). Adapter sequences were removed with Trim Galore. Reads were aligned with STAR to GRCh38/hg19 (human) or GRCm38/mm10 (mouse) and quantified to gene counts using RSEM. Gene expression values between samples were normalized using the geometric method in Cuffdiff. Analysis was performed using Pluto (https://pluto.bio). Gene ontology (GO) and pathway enrichment analyses were performed to unravel functional implications of the differentially expressed genes. Each GO term with a *p* value less than 1 × 10^−2^ was considered to be significantly enriched. Reverse transcription (RT)-PCR was performed as previously described ^59^.

### Enhancer RNA Analysis

Because enhancers are located in the gene body in genes tested, anti-sense eRNAs in enhancer regions were measured using quantitative RT-PCR.

### Neurosphere Culture

Neurospheres were generated from mouse cerebella (P5) through the meticulous process of cerebellum dissection, cell straining, and plating. Cells were plated at a density of 5 × 10^5^ cells per dish in a Neurobasal medium with B27 supplement, 20 ng/ml Epidermal Growth Factor (EGF), 20 ng/mL basic Fibroblast Growth Factor (bFGF), and 2 μg /mL heparin. Within three to five days, the cells self-organized into free-floating neurospheres.

### RNA Interference, *Kmt2d* Deletion, and Gene Expression in Neurospheres

siRNAs were used to conduct knockdown experiments (Supplementary Table 1). The neurospheres were transfected with siRNAs using Lipofectamine RNAiMAX. After a span of 48 hours, the cells were collected for RNA isolation.

Cerebellar neurospheres were placed in 60 mm dishes with a cell density of 2 × 10^4^ per dish. The deletion of *Kmt2d* in neurospheres was achieved using Adeno-Cre viruses at a rate of 1× 10^4^ pfu. For control, neurospheres infected with Adeno-Empty viruses were utilized. RNA was collected from the cells after 48 hours of infection with Adeno Empty or Adeno-Cre and analyzed using RT-PCR to confirm the deletion of *Kmt2d*.

### Measurement of Oxygen Consumption Rate

Cells were seeded in triplicate in a 12-well plate. Wells containing medium but no cells were used as baseline readings. Oxygen consumption was monitored as previously described ^10^, with some modifications. Briefly, cells (100 µg proteins) were incubated with the Metformin (50 mM) in buffer containing 250 mM sucrose, 30 mM K_2_HPO_4_, 1 mM EGTA, 5 mM MgCl_2_, 15 mM KCl, and 1 mg/ml bovine serum albumin supplemented with respiratory substrates and 50 nM MitoXpress, an oxygen-sensitive phosphorescent dye (LUXCEL, Cork, Ireland). On the second day, the medium in each well (including the wells without cells) was changed with 1 ml fresh medium. Rotenone (2 µM) and oligomycin A (1 µM) were used as 100% baseline for complex I and complex V inhibition, respectively. The areas under the curve were used for calculations. Oxygen consumption was then measured in real-time for 60 min at 37°C in 96-well plates using a spectrofluorimeter (Tecan Infinite 200; λExcitation 380nm; λEmission 650nm).

### ChIP Assays, ChIP-seq, and CUT&RUN

ChIP assay was performed using a modified version of the previously described method ^59,81^. Cerebellar tissues were processed for ChIP using Ren lab’s protocol from Roadmap Epigenome (https://www.encodeproject.org). The ChIP-seq libraries were prepared using NEB adapters as described earlier ^10^. Libraries were multiplexed together, and sequencing was performed in Hiseq2000 or HiSeq4000 (Illumina). CUT&RUN assays were performed using CUTANA ChIC/ CUT&RUN Kit. Analysis was performed with Pluto (https://pluto.bio) software.

### Inhibitor Experiments Using Cell Lines

AKP or *Ptch^+/−^* cell lines were cultured, and cells were seeded in four replicates in 96-well plates at a density of 1.5×10^3^ cells. The cells were treated with a range of concentrations of ORY-2001-Vafidemstat (0 to 1000 nM), Metformin (0.1 to 30 mM), SAHA (0 to 20 μM), TIC10/ONC201 (1 to 500 μM), SCH772984 (0 to 30 μM), Capivesertib (1 to 30 μM), Romidepsin (0 to 30 μM), and KDM5-C70 (0 to 30 μM) for 72 h (Supplementary Table 1). The cells were replenished with inhibitor-containing medium on every alternate day. Cell proliferation was quantified using Celigo followed by crystal violet staining. DMSO-treated cells were used as vehicle controls.

### Inhibitor Treatment of Xenograft Mouse Models

AKP or Ptch^+/−^ cell lines were cultured, and cells (5 × 10^6^) in 100 μl of Matrigel were subcutaneously injected in the both flanks of 6-to 8-week-old athymic nu/nu mice. After 8 days when the tumors were noticeable, the mice with tumors were randomly divided into 4 groups. Mice were treated with intraperitoneal injections of ORY-2001 (500 mg/kg body weight), Metformin (500 mg/kg body weight), or a combination of ORY-2001 plus Metformin on an alternate day until 16^th^ days (after cell injection). The drugs were prepared in a neurobasal medium with growth factors and Matrigel. The same medium was used for vehicle control. The tumors were measured every other day using a caliper. To calculate the tumor volume, the ellipsoid volume formula (1/2 x length x width x height) was used, as previously described ^10^. Upon completing their treatment, the mice were euthanized, and their tumors were collected for subsequent examination via histological analysis.

### Statistical Analysis

Prism GraphPad was mainly used for conducting statistical analyses. The mean and standard error of the mean (SEM) were presented from a minimum of three independent experiments. The two-tailed Student t-test was used to test statistical significance, unless stated otherwise. * (p < 0.05), ** (p < 0.01), and *** (p < 0.001) denote statistically significant differences. A two-sided log-rank test was used to determine statistical difference between tumor-free curves of different mouse models and between their survival curves. Such curves were generated using the Kaplan-Meier method.

---

**Extended Data Fig. 1.**
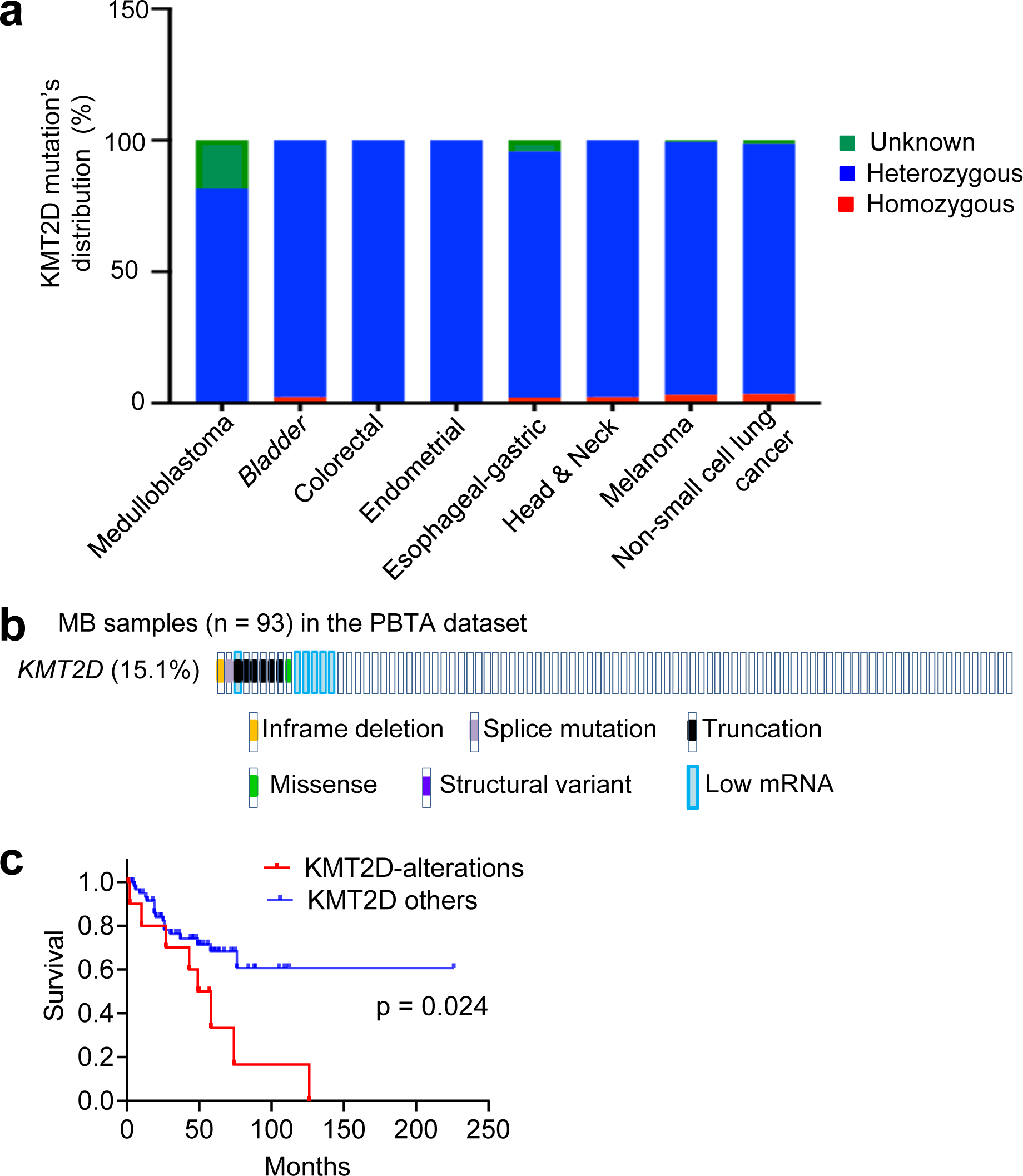
*KMT2D* is altered in medulloblastoma and other cancer, related to Figure 1: (**<u>a</u>**) High percentage (81-99%) of heterozygous mutations of the *KMT2D* gene occurred in MB and other types of cancer. The graph shows mutation status (unknown, heterozygous, and homozygous) in the *KMT2D* gene. Data (except MB data) were generated using MSK-IMPACT dataset in PanCancer studies in cBioPortal. Heterozygous states were determined using mutation’s allele frequency (0 – 0.5) in the dataset (**<u>b</u> & <u>c</u>**) Analysis of the PBTA MB dataset in the PedcBioportal database showed that KMT2D alterations (*KMT2D*-low/mutations) correlated with poor survival of MB patients. Low *KMT2D* mRNA levels denote those with Z score with ≤ 1.5.

**Extended Data Fig. 2.**
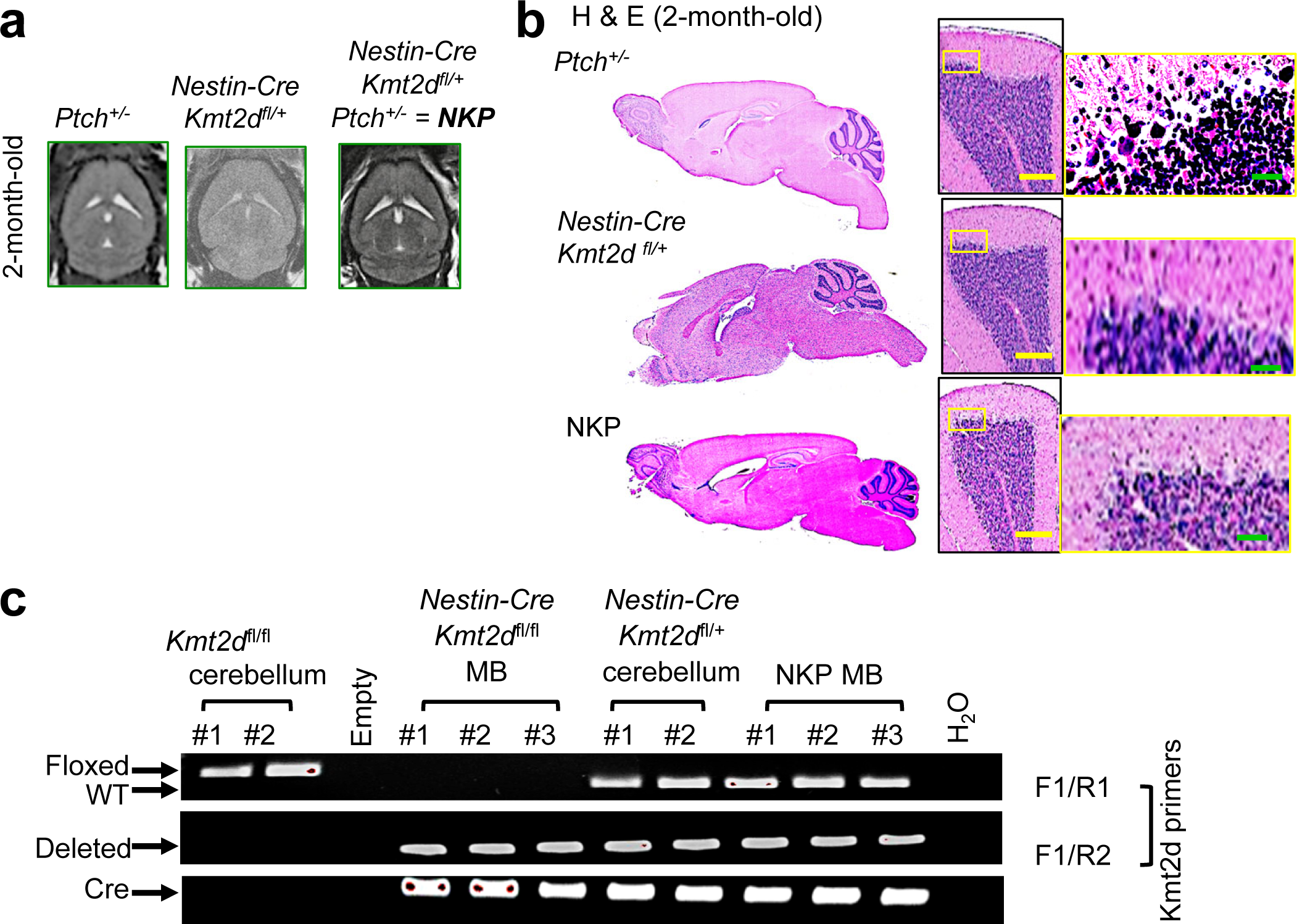
*Nestin-Cre*-mediated heterozygous loss of *Kmt2d* induced no obvious MB in the 2-month-old cerebellum, related to Figure 1: (**<u>a & b</u>**) MRI (**a**) and H & E staining (**b**) showed that *Nestin-Cre*-mediated heterozygous loss of *Kmt2d* induced no obvious sign of MB in the cerebellum in 2-month-old *Ptch^+/−^* mice. (**<u>c</u>**) Genotyping results using specific primers confirmed heterozygous deletion of *Kmt2d* in NKP MB. For comparison, normal *Kmt2d*^fl/fl^ cerebellum were used.

**Extended Data Fig. 3:**
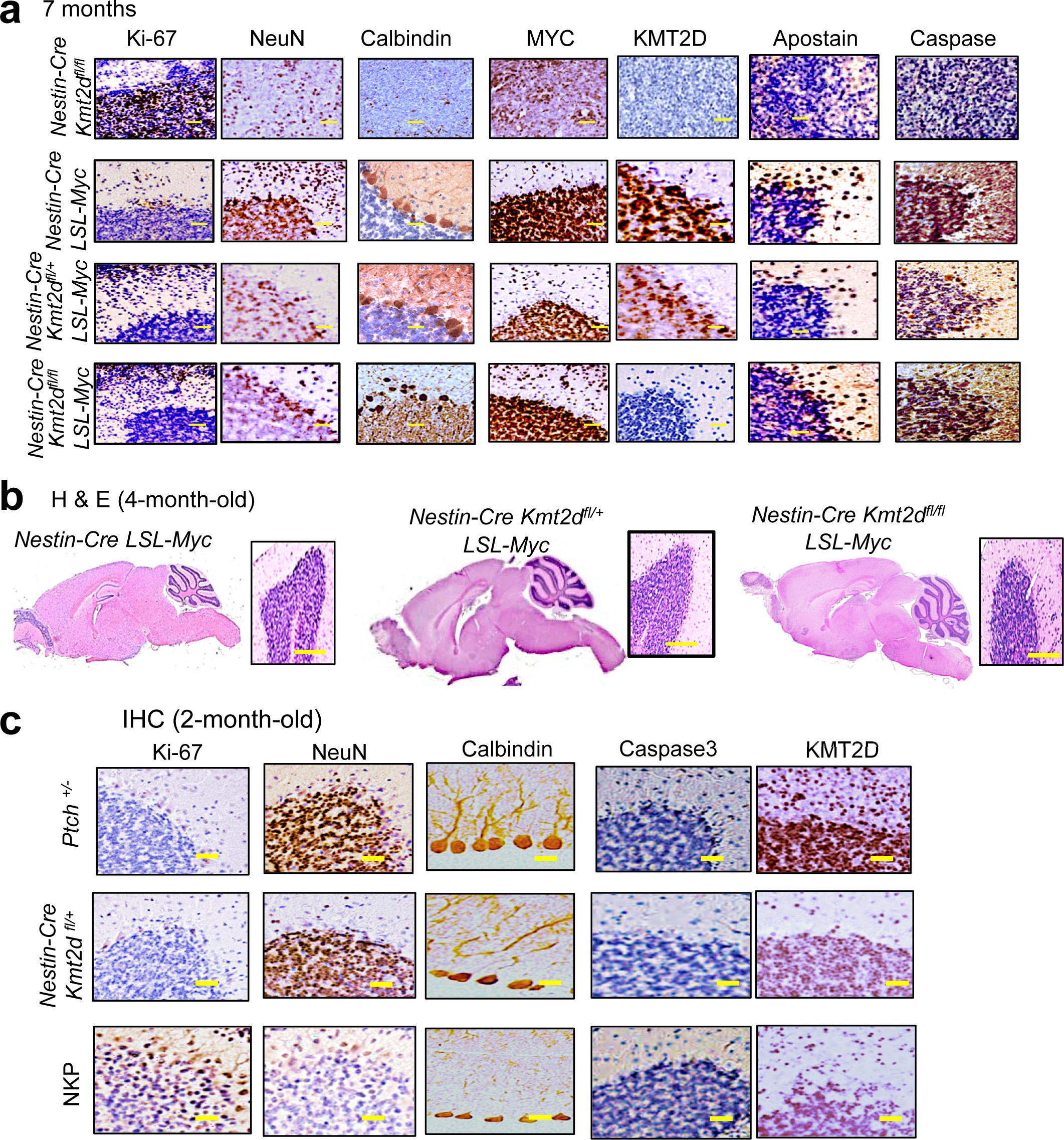
*Kmt2d* loss did not cooperate with *MYC* overexpression for MB genesis (a & b), whereas *Nestin-Cre*-mediated heterozygous loss of *Kmt2d* increased cell proliferation and negatively affected the neuronal marks in *Ptch*^+/−^ cerebellum (c), related to Figure 1. (**<u>a</u> & <u>b</u>**) IHC results showed that MYC overexpression increased the staining of cleaved Caspase and Apostain and (**a**). H&E staining did not show any obvious sign of MB in *Nestin-Cre LSL-Myc, Nestin-Cre Kmt2d^fl/+^LSL-Myc,* and *Nestin-Cre Kmt2d^fl/fl^ LSL-Myc (*4-month-old) mice (**b**). (**<u>c</u>**) IHC results showed that *Nestin-Cre*-mediated heterozygous loss of *Kmt2d* significantly increased Ki-67 signals (a proliferation marker) and reduced NeuN and Calbindin (neuronal markers) staining in *Ptch*^+/−^ cerebella (2-month-old mice). Scale bars, 100 µm.

**Extended Data Fig. 4:**
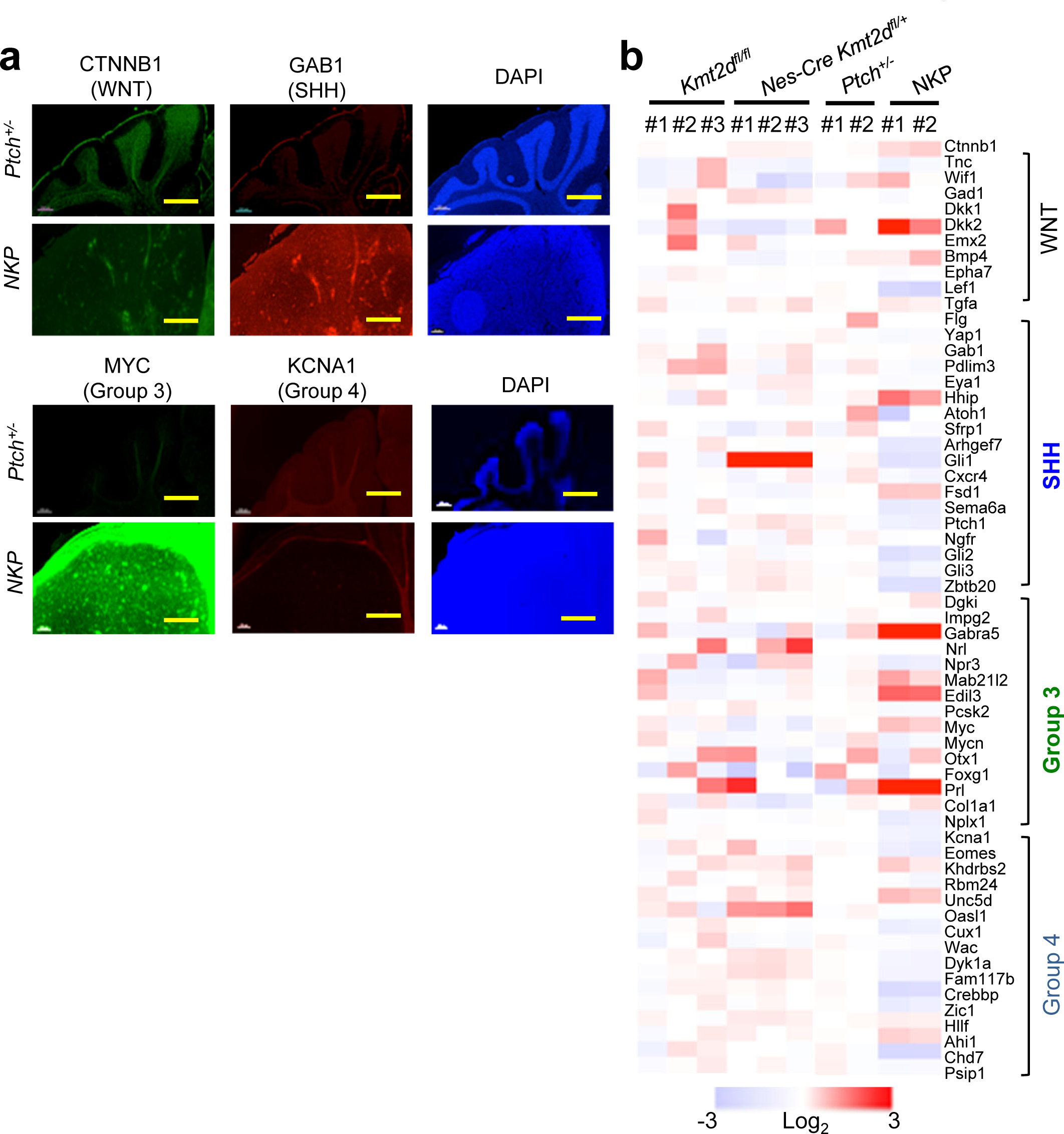
NKP MBs express markers for SHH MB and group 3 MB according to transcriptomic and IF analysis, related to Figure 1. (**<u>a</u>**) Immunofluorescence (IF) staining of *Ptch*^+/−^ and NKP tumors showed that signals of GAB1 (SHH group marker) and MYC (Group 3 marker) were increased in 4-month-old NKP MB compared with *Ptch*^+/−^ and *Kmt2d^fl/fl^* cerebella. However, no obvious difference was observed for CTNNB1 (WNT group marker) and KCNA1 (Group 4 marker) signals between 4-month-old NKP MB and control (*Ptch*^+/−^) cerebella. Scale bars, 200 µm. (**<u>b</u>**) Expression levels of some signature genes for Group 3 and SHH MB were elevated in NKP tumors compared with control (*Ptch^+/−^*). For direct comparison, we added previously published expression data for *Kmt2d^fl/fl^* and *Nes-Cre Kmt2d^fl/+^* cerebella.

**Extended Data Fig. 5:**
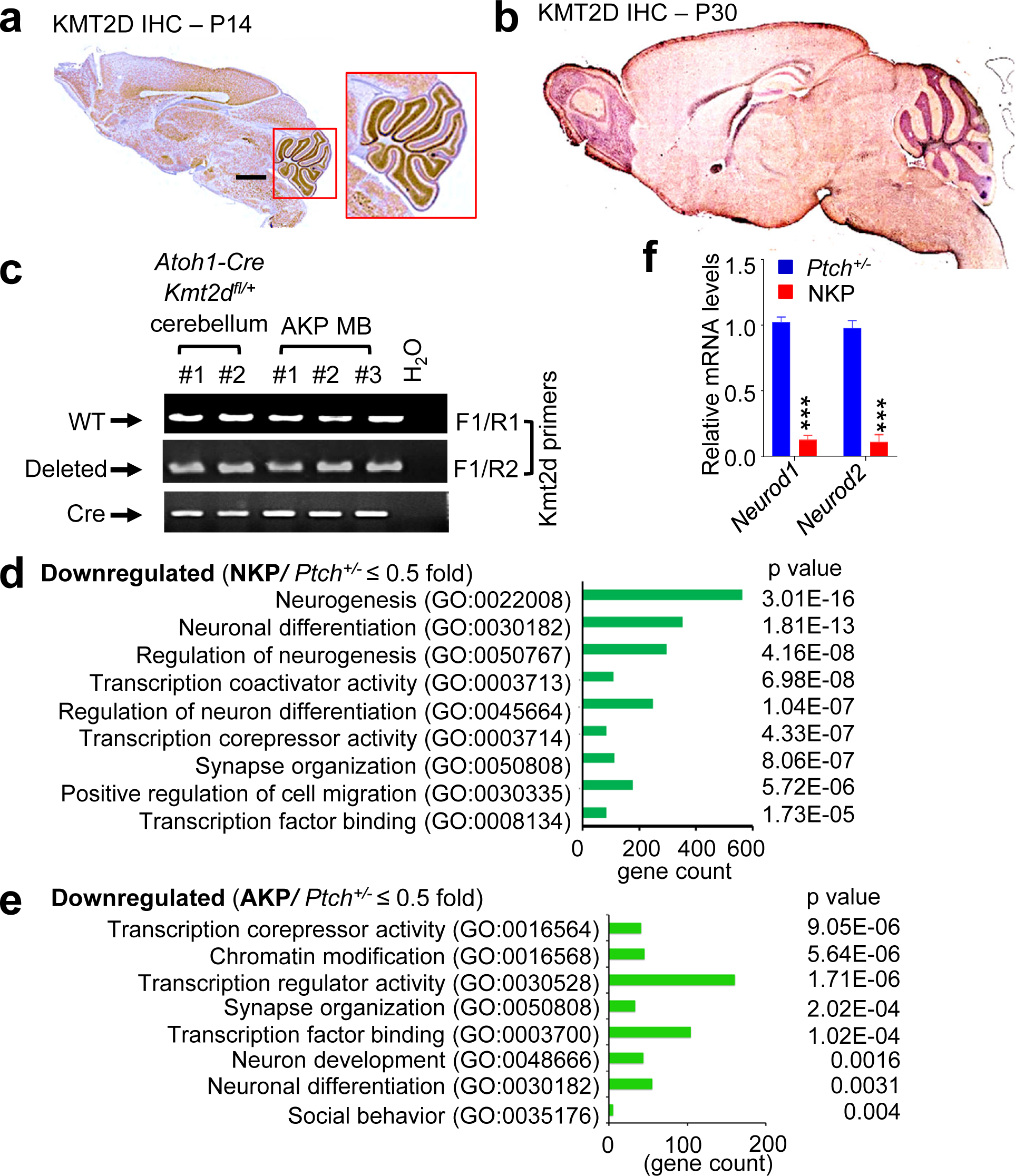
KMT2D is highly expressed in the cerebellum, and heterozygous *Kmt2d* loss downregulates neuronal genes and transcriptional corepressor genes, related to Figure 2 & 3. (**<u>a</u> & <u>b</u>**) IHC staining of normal mouse brain showed high KMT2D signals in granular cell layers in the cerebellum at postnatal day 14 and 30 (P14 & P30) (**<u>c</u>**) Genotyping results using specific primers confirmed heterozygous deletion of *Kmt2d* in AKP MB. (**<u>d</u> & <u>e</u>**) Gene ontology analysis showed gene expression programs downregulated by heterozygous *Kmt2d* loss. NKP (**d**) or AKP (**e**) cerebellar tissues were compared to *Ptch*^+/−^ cerebellar tissues (i.e., NKP */Ptch*^+/−^ and AKP/*Ptch*^+/−^). GO pathways were determined using the Panther analysis. (**<u>f</u>**) Quantitative RT-PCR analysis showed that *Neurod1* and *Neurod2* were downregulated in NKP cerebella as compared to *Ptch*^+/−^ cerebella.

**Extended Data Fig. 6:**
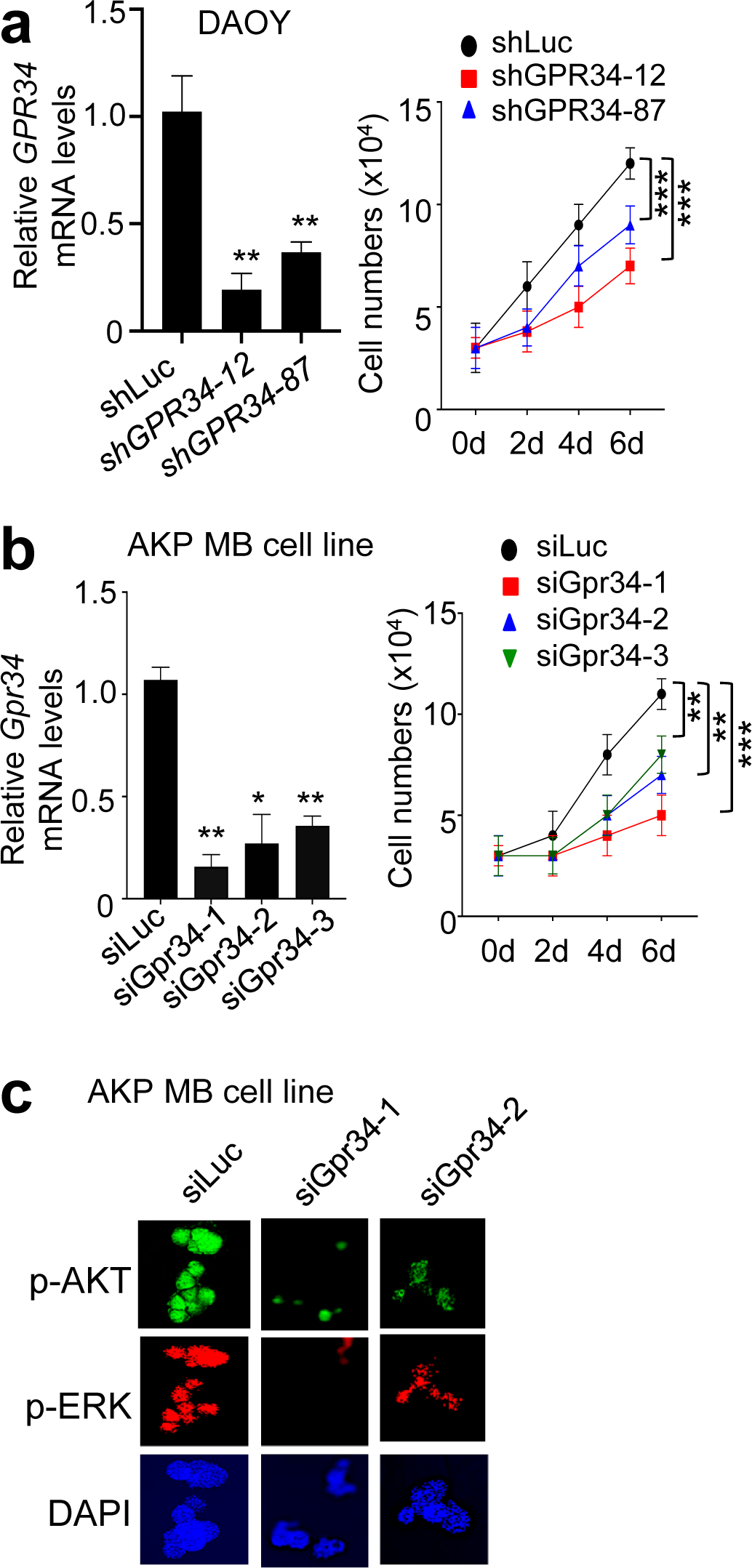
Knockdown of GPR34 decreased the proliferation of human DAOY MB cells and mouse AKP MB cells, related to Figure 4. (**<u>a</u> & <u>b</u>**) Knockdown of GPR34 decreased the proliferation of human DAOY MB (**a**) and mouse AKP MB (**b**) cell line. (**<u>c</u>**) IF staining showed that GPR34 knockdown reduced p-AKT and p-ERK signals in AKP MB cell line. DAPI was used for nuclear staining. Statistical analysis was performed using two-tailed Student’s t test. Data are represented as the mean ± SEM. *p < 0.05, **p < 0.01 and ***p < 0.001.

**Extended Data Fig. 7:**
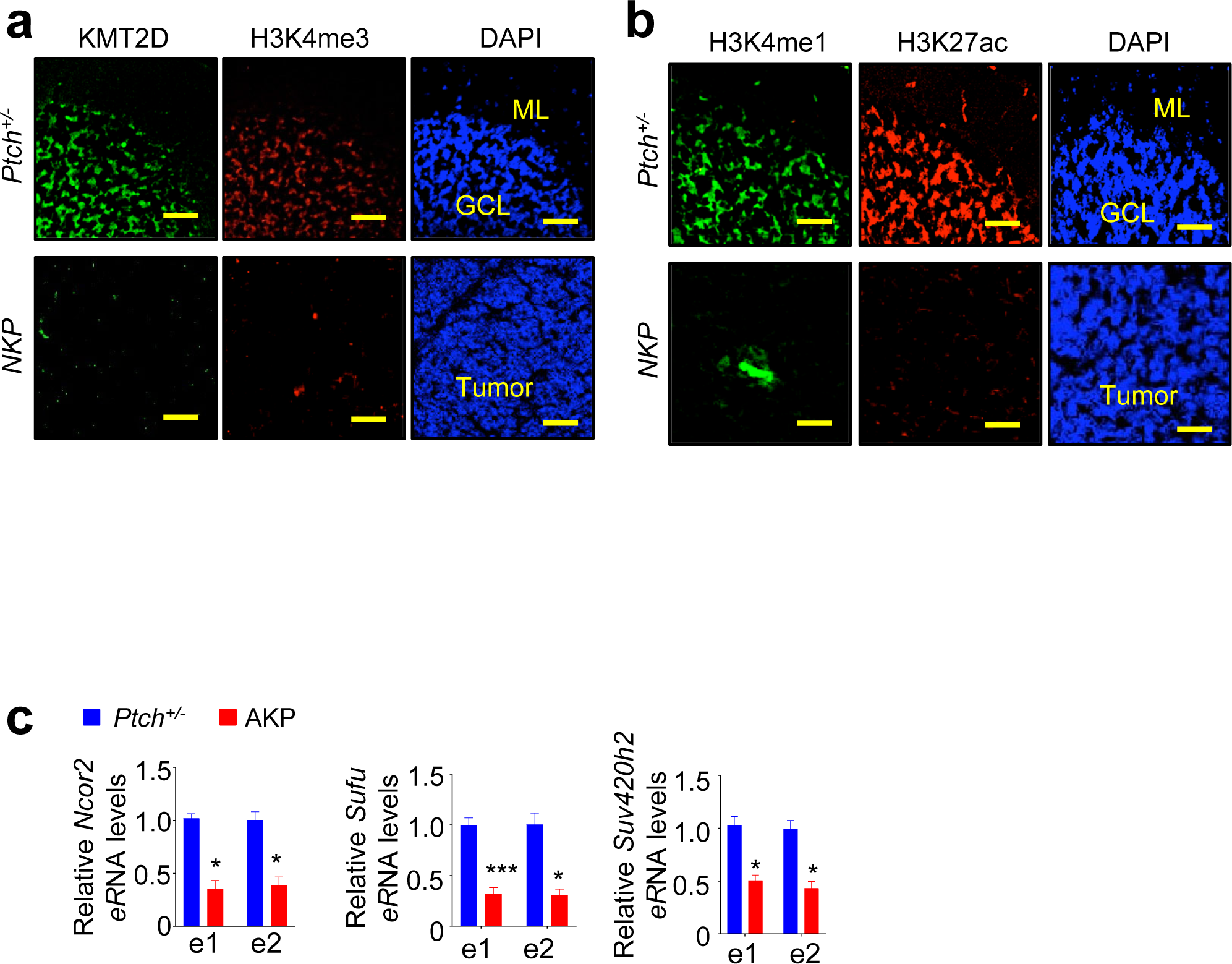
*Nestin-Cre*-mediated heterozygous loss highly reduced the epigenomic marks H3K4me3, H3K4me1 and H3K27ac, and *Atoh1-Cre*-mediated heterozygous loss of *Kmt2d* reduced eRNA levels in *Ncor2*, *Sufu*, and *Suv420h2.* related to Figure 7. (**<u>a</u> & <u>b</u>**) Immunofluorescence (IF) analysis for KMT2D, H3K4me3, H3K4me1, and H3K27ac were performed using NKP and *Ptch^+/−^* cerebellum. Scale bars, 50 µm. (**<u>c</u>**) Quantitative RT-PCR showed that *Atoh1-Cre*-mediated heterozygous loss of *Kmt2d* decreased eRNA levels in *Ncor2*, *Sufu*, and *Suv420h2* enhancers. *See* Fig. 7G for the locations of eRNA amplicons.

**Extended Data Fig. 8:**
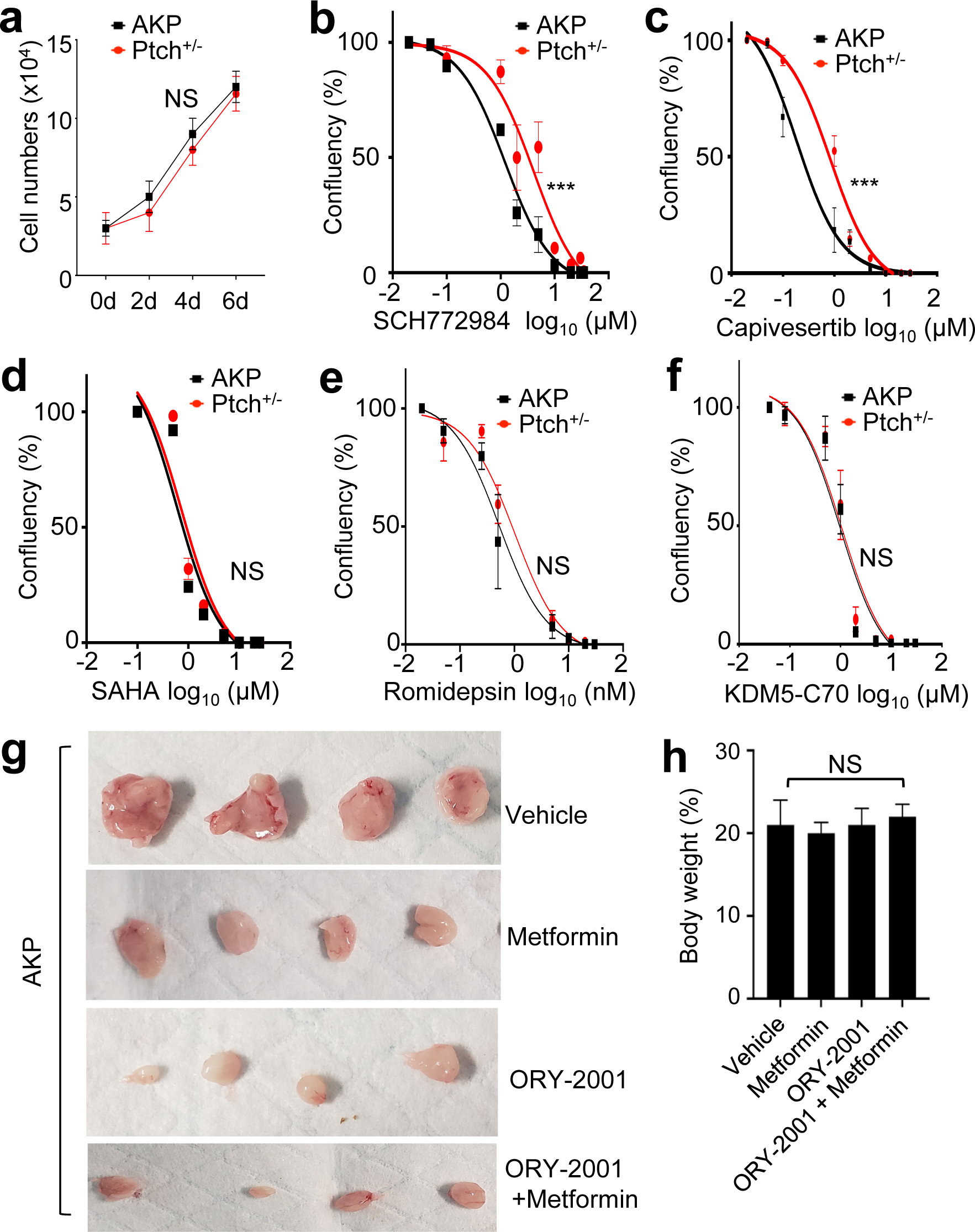
The effects of different inhibitors on the proliferation of AKP and *Ptch^+/−^* MB cell line, related to Figure 8. (**<u>a</u>**) AKP and *Ptch^+/−^* MB cell lines showed similar cell proliferation rate. (**<u>b</u>‒<u>f</u>**) Dose response experiments showed that SCH772984 and Capivesertib preferentially inhibited the proliferation of AKP cell line compared to *Ptch^+/−^* MB cell line whereas SAHA, Romidepsin, and KDM5-C70 did not. Cells were treated with different concentration of inhibitors for 72h. Data are represented as the mean ± SEM. NS, not significant; ***p < 0.001. (**<u>g</u> & <u>h</u>**) The combination of Metformin and ORY-2001 synergistically inhibited tumorigenic growth of AKP MB cells in a mouse subcutaneous xenograft model (**g**) but did not affect mouse weight (**h**). Tumors were dissected from the mice (**g**). NS, not significant.

## Notes

https://sciencecast.org/casts/3fe2q5snd4kw?t=jQF73lUSriZGQdwd%2B6mRp8ckbcc6tKVyKSwwd7Yl6wDyGVstIBBbXVvp%2FIIQsbggQknzlLpxv7nwg4g18UsikQ%3D%3D

### Summary of Updates

Revised text and supplementary table

